# Neural decoding of speech using deep neural ensembles

**DOI:** 10.64898/2026.06.02.729705

**Authors:** Seonghyun Yoon, Donald T. Avansino, Sasidhar Madugula, Alisa D. Levin, Chaofei Fan, Benyamin Abramovich Krasa, Akansha Singh, Christina Vo, Nick V. Hahn, Nicholas S. Card, Zachery Fogg, Maitreyee Wairagkar, Samuel R. Nason-Tomaszewski, Brandon G. Jacques, Payton H. Bechefsky, Carrina Iacobacci, Darrel R. Deo, Leigh R. Hochberg, David M. Brandman, Sergey D. Stavisky, Nicholas Au Yong, Chethan Pandarinath, Jaimie M. Henderson, Francis R. Willett

## Abstract

Speech brain-computer interfaces (BCIs) can restore rapid communication to people with paralysis, but decoding errors still limit performance. In recent brain-to-text decoding competitions, deep ensemble methods, which combine predictions from multiple independently trained decoders, have delivered striking accuracy improvements and account for the largest gains over baseline approaches. However, these methods have not previously been tested in real-time, require substantial computational resources, and their performance under various clinically relevant constraints remains poorly understood. Here, we present the first closed-loop test of deep ensembles in a participant with bilateral intracortical microelectrode arrays, demonstrating a reduction in word error rate from 33.7% to 26.0% on a large-vocabulary task. Using additional data from three participants, we then assess how these gains depend on baseline error rate, training dataset size, and ensemble size, including the resource-accuracy tradeoffs most relevant for real-world deployment. Finally, we introduce a computationally efficient pseudoensembling approach based on test-time augmentation that improves decoding accuracy while requiring only a single base decoder, greatly reducing the computational burden of ensembling. Together, these results show that the benefits of deep ensembling can be realized in real time and under practical resource constraints, bringing speech BCIs closer to broader clinical adoption.

## Introduction

Speech brain-computer interfaces (BCI) have sought to restore rapid and highly accurate communication to people with paralysis by translating neural signals evoked by imagined or attempted speech into text^1–7^. In one study, an individual with amyotrophic lateral sclerosis (ALS) maintained independent use of an intracortical speech BCI for over 3,800 cumulative hours across 600 days at 56.1 words per minute, demonstrating their potential to restore fast and reliable communication in individuals with paralysis^5^.

While prior results are indeed impressive, decoding performance can vary due to individual differences in the degree and etiology of paralysis^6–8^, yield and cortical coverage of the implanted arrays^3,4,7,9,10^, vocabulary size^1–3,7^, speech mode^11–13^, and available training data^2,3,7,14^. The resulting variations in word error rate (WER) can substantially impede communication (Fig. 1b) for a few reasons. First, natural language follows Zipf’s law, in which a small number of common, generic words account for the majority of word occurrences, while a large number of rare words convey specific meaning^15,16^. Errors on semantically salient words may be infrequent, but could cause a disproportionately large shift in the perceived topic of a sentence or even invert its intended meaning^17^. Furthermore, prior work in automatic speech recognition has shown that modest increases in error rate can increase user correction overhead, disproportionately reducing effective communication rate^18^. For people with severe speech and motor disorders, the failure to reliably communicate using assistive technologies may result in frustration^19^, user fatigue^19^, social isolation^20^, hinder effective medical care^21^, and even influence decisions about life-sustaining treatment^22,23^. Consequently, improving decoding accuracy remains a central objective for advancing the clinical viability of speech BCIs. WERs of 25% and 5% have respectively been suggested as minimally viable and reliably useful system performance targets^16^.

**Fig. 1.**
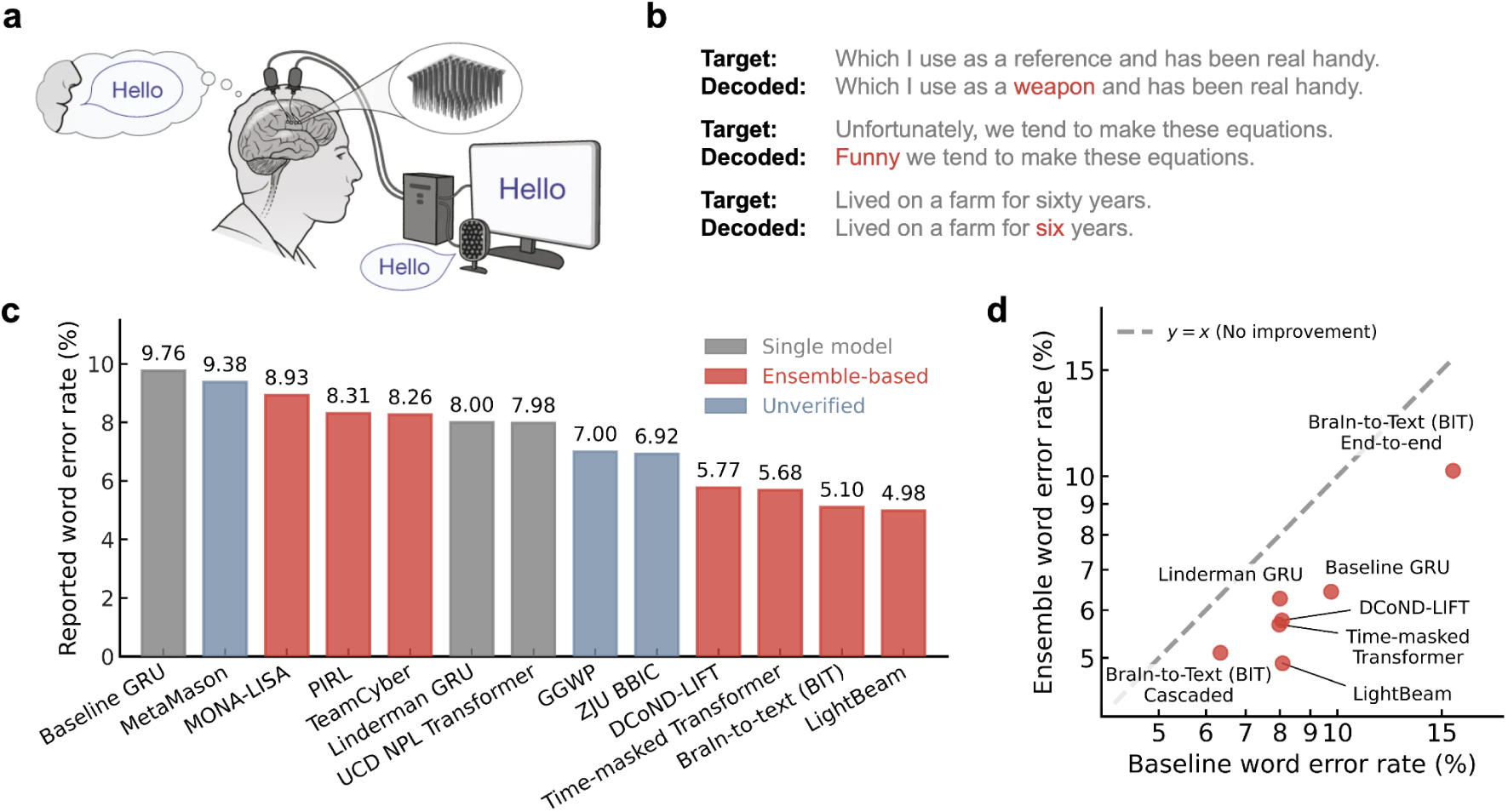
Survey of recent advances in brain-to-text decoding. **a,** A speech brain-computer interface can translate a user’s neural activity into intended speech. **b,** Example outputs from a brain-to-text decoder illustrating that modest, single-word errors can still substantially alter semantic meaning. The decoder achieved an 8.10% word error rate on the Brain-to-Text ‘24 Benchmark. Nevertheless, the intended meaning is altered through changes in the described object (“reference” to “weapon”), speaker stance (“unfortunately” to “funny”), and quantitative content (“sixty” to “six”). Example sentences were selected from a separate held-out partition. **c,** Deep ensembles were commonly used among the top-performing entries of the Brain-to-Text ‘24 Benchmark. Of the nine verified entries that outperformed the baseline models, seven methods used deep ensembles. **d,** For the participant T12 from the Brain-to-Text ‘24 benchmark, deep ensembles improved performance across a range of model architectures and word error rates. Only ensembles using ten base decoders are shown to avoid ensemble size as a confounding factor.

To catalyze collaborative efforts in improving speech BCI performance, we launched two online competitions, the Brain-to-Text ‘24 and ‘25 Benchmarks, with open intracortical datasets collected from two BrainGate2 clinical trial participants^24,25^. More than 500 entries from the machine learning and neuroscience communities have yielded substantial insights into effective decoding strategies. One key finding was that the highest performers consistently employed deep ensembles (Fig. 1c)^5,26–31^, a technique in deep learning that combines predictions from multiple independently trained models^32^. Prior works in traditional sequence transduction domains such as automatic speech recognition have widely adopted deep ensembling due to its strong performance gains and ease of integration into existing pipelines^33–35^. In offline studies decoding speech from intracortical neural activity, Benster et al. first demonstrated the efficacy of aggregating ensemble-generated hypotheses using a large language model (LLM)^27^. Li et al. further employed large-scale ensembles and LLM fine-tuning to improve hypothesis generation and selection, substantially reducing the WER on the Brain-to-Text ‘24 benchmark from 9.76% to 5.77%^28^. More recently, deep ensembles have been shown to be effective across different base model architectures and performance regimes for the single-participant dataset released in the Brain-to-Text ‘24 benchmark (Fig. 1d)^26,28,29,31,36^, suggesting its potential relevance even as architectural advances continue to improve single-model performance.

Despite these promising offline results, deep ensembles have not been tested in real-time settings. As a result, it is unclear to what extent these strong offline accuracy gains can translate into real-time performance. Although well-evaluated offline results can be informative in decoder development, they may fail to reflect possible distribution shifts^3–5^, error perception^37^, and visual feedback^10,38^ present in neural signals during closed-loop evaluation. In addition, offline results can overfit to the particular evaluation dataset, as hyperparameters can be optimized with knowledge of the resulting test set performance. In contrast, real-time testing can provide a stronger standard of evidence^39^, since evaluation is performed only once after the system design is fixed, meaning performance cannot arise from overoptimizing to a specific dataset. Beyond decoding performance, system-level considerations critical for the real-world deployment of deep ensembles also remain unexplored, such as inference latency, computational constraints, and the cloud-based system architectures that may be able to support the computational cost of real-time ensemble inference.

To bridge this gap, we demonstrate that a deep ensemble speech BCI using cloud computational resources can substantially improve accuracy in a real-time, closed-loop paradigm. Furthermore, using a large dataset of neural activity during attempted speech (67.7 hours, 3 participants, 4 array configurations), we systematically characterize deep ensemble performance with respect to considerations relevant to clinical translation, including diversity in baseline error rates, amount of training data, and computational resources. Finally, we also propose a lightweight “pseudoensembling” solution for resource-constrained deployment settings, which uses stochastic perturbations during inference to generate a hypothesis set, bypassing the computational overhead of full ensembling.

## Results

### Deep neural ensembles improve real-time decoding accuracy

To evaluate the real-time performance of deep ensembles, we implemented and tested an ensemble-based speech BCI with one BrainGate2 clinical trial participant (T12). Neural activity was recorded from six intracortical microelectrode arrays placed bilaterally along the precentral gyri targeting areas related to speech production, with two arrays each in right area 6v, right area 55b, and left area 6v (Extended Fig. 1). The participant performed a copy-speaking task in which she attempted to speak prompted sentences drawn from a 125,000-word vocabulary.

Our ensemble speech BCI consisted of ten base decoders independently trained from randomly initialized weights (Fig. 2a). Base decoders used a RNN-based architecture which had been extensively validated in multiple high-performing real-time speech BCI studies^3,4,6,7^ (see Table 4 for hyperparameters). As in prior works^4,5,40^, each base RNN was also continuously recalibrated during inference to adapt to nonstationarities. To provide closed-loop feedback while T12 attempted speech, one base decoder was chosen at random to update and display its current prediction every 80 milliseconds (ms). After T12 indicated the end of a sentence with a button press, each base decoder computed its hypothesis in parallel, and a “merger” LLM produced a coherent final prediction. The merger LLM was fine-tuned ahead of time to best combine the ensemble-generated candidate predictions.

**Fig. 2.**
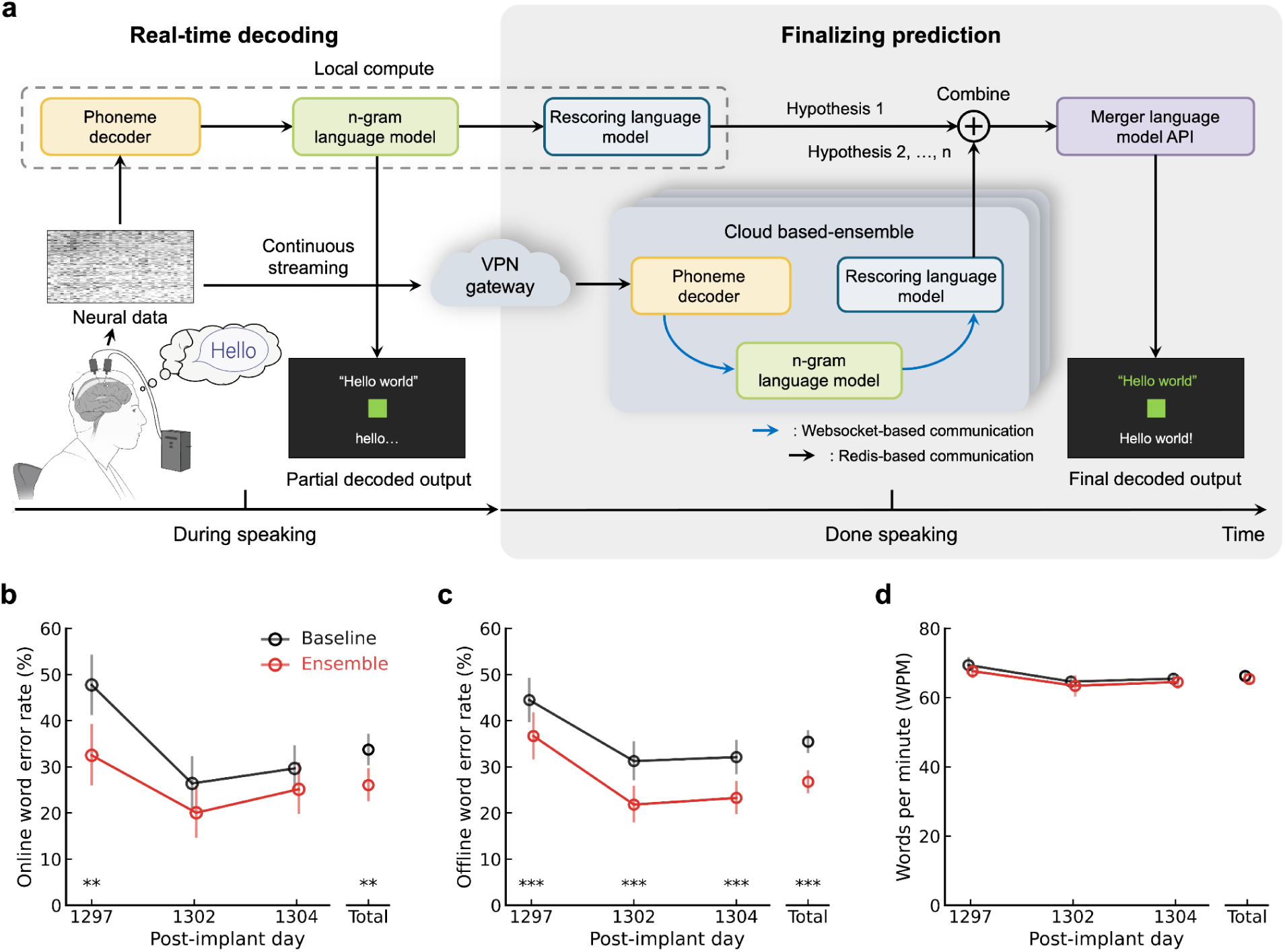
Implementation and real-time performance of the ensemble speech BCI. **a,** System diagram of the ensemble speech BCI. Each base decoder consisted of a recurrent neural network that maps neural activity to phoneme probabilities, and two language models that infer the most likely word sequence. During speech, one locally deployed base decoder produces a partial prediction. Concurrently, neural data is streamed to the cloud platform via a secure VPN gateway. After the user indicated the end of a sentence, each hypothesis was computed in parallel on cloud, and aggregated via a fine-tuned merger LLM. The displayed output is subsequently updated. **b,** Online word error rates of the baseline and ensemble BCI. Comparisons were conducted via paired permutation test (*n*=10,000, * if *p* < 0.05, ** if *p* < 0.01, *** if *p* < 0.001). Error bars indicate 95% CIs computed via bootstrapping (*n*=10,000). **c,** Same as **b,** but for an offline “replay” of the session to compare performance on identical sentences. **d,** Same as **b,** but for speech rate (words per minute).

The base decoders were distributed across a network of 20 cloud virtual machines, to which neural features were continuously streamed over a secure VPN gateway. One base decoder used for displaying partial predictions was deployed locally to minimize feedback latency, incurring negligible delay beyond an intrinsic delay of 80 ms due to the RNN’s update window size. We note that this decoder could also be deployed on the cloud to enable a fully cloud-based BCI, but with an added feedback latency of 273 ms (95% CI, 265–282 ms; averaged across 10 sentences) during a pre-session network test.

Finally, the merger LLM was fine-tuned on a held-out set during decoder training (3,162 sentences) and accessed via a remote API. To maximize real-time performance, the final prediction was selected as the highest-probability output after performing parallel inference of three fine-tuning checkpoints (similar to checkpoint ensembling^41^). Where local deployment is necessary, this method could be implemented in a memory-efficient manner by changing only the low-rank adapters while re-using the base model^42^.

We compared our ensemble speech BCI to a single-decoder baseline in interleaved blocks of 40 sentences, yielding a total of 14 evaluation blocks over three session days (Supplementary Video 1). We found that our ensemble BCI substantially reduced the WER from 33.7% to 26.0% (*p* < 0.01; Fig. 2b) while T12 maintained a speech rate of 65.4 words per minute (Fig. 2d).

Because the interleaved design necessarily evaluated the ensemble and baseline on different sentences, we replicated the decoder states offline to compare performance on identical sentences. Consistent with online results, ensembling significantly reduced WER from 35.5% to 26.8% (*p* < 0.0001; Fig. 2c). We also evaluated performance offline without using multiple merger LLM checkpoints. Aggregating hypotheses with just a single merger LLM checkpoint instead of choosing the best out of three predictions (as performed during real-time evaluation) still reduced WER to 29.2% (*p* < 0.0001), suggesting checkpoint ensembling provided additional gains but was not necessary to improve accuracy.

### Deployment-relevant considerations for ensemble-based BCIs

We next characterized the latency introduced by ensemble decoding. During real-time evaluation sessions, ensembling incurred an extra delay of 5.25 seconds to display the final prediction on average (Fig. 3a). Step-wise latency decomposition (Fig. 3b) showed that the merger LLM API had the largest contribution of 36.9% (mean 2.77 seconds; 95% CI, 2.50–3.06 seconds). This was followed by completion of neural data readout (34.2%; mean 2.57 seconds; 95% CI, 2.26–2.91 seconds), phoneme decoding (16.7%; mean 1.25 seconds; 95% CI, 1.21–1.29 seconds), and language model decoding (11.8%; mean 887 ms; 95% CI, 841–935 ms).

**Fig. 3.**
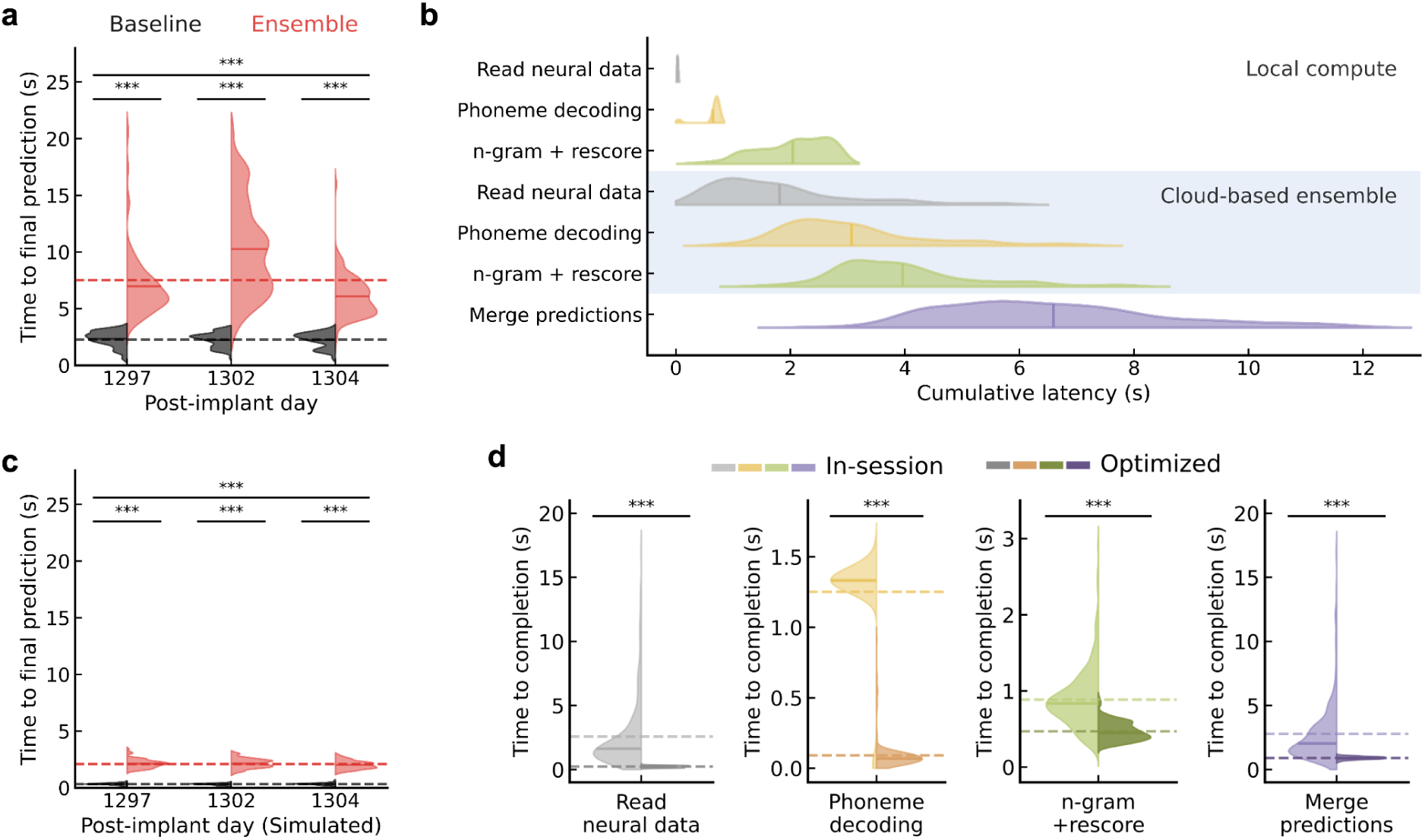
In-session and optimized decoding latency of the ensemble speech BCI. **a,** Time taken to produce the final prediction after the participant signified her end of speech. Solid and dotted horizontal lines represent per-session and global average times, respectively. Comparisons were performed via paired permutation test (*n*=10,000, *** if *p* < 0.001). **b,** Cumulative completion times of each decoding stage during a typical trial, shown for the 90th percentile of trials by end-to-end latency. For cloud-based stages, completion time was defined as the slowest base decoder on each trial. Vertical lines for each step represent the mean cumulative time taken until completion. **c,** Same as **a**, but for the optimized baseline and ensemble BCIs during a replayed simulation of the real-time evaluation sessions. The y-axis range matches **a** for a direct comparison. The replay was purposefully evaluated under more suboptimal network conditions than those during the real-time sessions. **d,** Completion times for each decoding stage before and after optimization. These results yielded key principles for meeting clinically relevant latency levels.

Since our initial implementation was intended as a proof-of-concept to primarily test the accuracy gains of ensembling, we then evaluated whether ensemble BCIs could be optimized to meet clinically practical response times. In an offline simulation replaying real-time evaluation session data, our optimized system achieved an end-to-end delay of 2.09 seconds (95% CI, 2.04–2.14 seconds). This was only 1.76 seconds slower than the optimized baseline and 5.43 seconds faster than our previous ensemble BCI implementation (Fig. 3c).

Implementation details can be found in Extended Fig. 2a for a system architecture and Extended Figs. 3 and 4 for operational finite-state machines. Hardware used for both real-time evaluation and offline simulation can be found in Table 1. However, we note that this should not be interpreted as a minimum requirement as hardware utilization was significantly lower (Table 2).

**Fig. 4.**
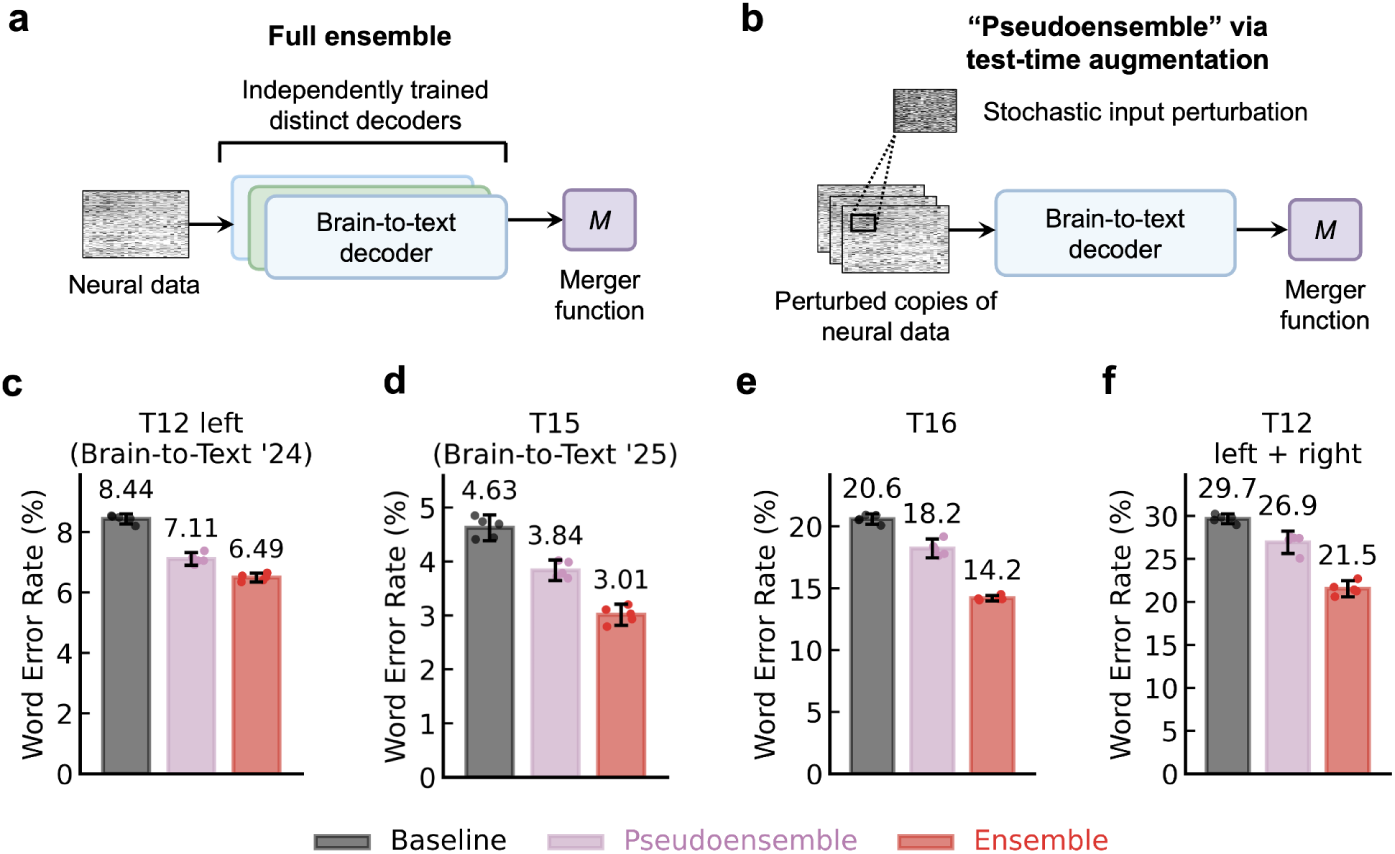
Effectiveness of deep ensemble methods across participants, array configurations and performance regimes. **a,** Abstracted diagram of a full ensemble decoder. **b,** Abstracted diagram of a pseudoensembling decoder. Stochastic input perturbations produce diverse predictions, approximating a full ensemble with only a single base decoder. **c,** A comparison of ensemble-based methods with the single-decoder baseline for T12 left hemisphere data. Bar heights denote the average word error rate across five training seeds, each overlaid as a point. Error bars show 95% CIs. **d–f,** Same as **c,** but for T15, T16, and T12 left and right hemisphere data, respectively.

**Table 1.** Hardware provisioned for the ensemble speech BCI.

| Purpose of compute node | Hosted on | CPU specification | Memory size | GPU specification | Qty |
| --- | --- | --- | --- | --- | --- |
| Neural data processing and Redis server | Local rig | Intel i9-12900K (3.20 GHz, 16 cores, 32 threads) | 128 GB | N/A | 1 |
| RNN inference and recalibration, display | Local rig | AMD Ryzen 9 5950X (3.40 GHz, 16 cores, 32 threads) | 128 GB | NVIDIA RTX 4090 (24 GB VRAM) | 1 |
| 5-gram language model and rescoring LLM | Local rig | AMD Ryzen Threadripper Pro 9995WX (2.50 GHz, 24 cores, 48 threads) | 512 GB | NVIDIA A6000 (48 GB VRAM) | 1 |
| VPN server and data relay | Google cloud | Intel Xeon E5-2699 v4 (2.20 GHz, 2 cores, 4 threads) | 16 GB | N/A | 1 |
| RNN inference and recalibration | Google cloud | Intel Xeon Platinum 8273CL (2.20 GHz, 16 cores, 32 threads) | 128 GB | NVIDIA L4 (24 GB VRAM) | 9 |
| 5-gram language model | Google cloud | AMD EPYC 7B13 (2.45 GHz, 56 cores, 112 threads) | 224 GB | N/A | 1 |
| Rescoring LLM | Google cloud | Intel Xeon Platinum 8273CL (2.20 GHz, 4 cores, 8 threads) | 32 GB | NVIDIA L4 (24 GB VRAM) | 9 |

**Table 2.** Peak hardware utilization of cloud virtual machines used for the optimized ensemble speech BCI.

| Purpose of compute node | Equivalent vCPUs used during peak utilization trial | Peak memory | Peak GPU memory |
| --- | --- | --- | --- |
| VPN server and data relay | 0.32 | 0.039 GiB | N/A |
| RNN inference and recalibration | 1.23 | 10.92 GiB | 18.22 GiB |
| 5-gram language model | 7.45 | 9.98 GiB | N/A |
| Rescoring LLM | 1.13 | 1.61 GiB | 13.98 GiB |

From this result, we identified three key principles that may guide future ensemble speech BCI designs. First, because intracortical neural data are high-rate and multichannel, redundant local-cloud communication can rapidly increase network overhead. Instead of streaming data directly to each base decoder and online recalibration process, we designed a single cloud relay node that receives neural data from the local rig, and distributes to individual base decoder machines via the high-bandwidth intra-cloud network. This relay architecture reduced neural data readout time by 91.2% (mean 228 ms; 95% CIs, 217–238 ms). Second, because brain-to-text decoding applies a fixed decoding pipeline to each new neural data window, pre-computation can avoid recurrent overhead. Caching key-value tensors of the merger LLM’s fixed prompt prefix, together with more efficient cache management^43^, improved hypothesis aggregation time by 67.4% (mean 903 ms; 95% CI, 889–916 ms) when using three Qwen3-4B^44^ fine-tuning checkpoints on a local GPU (NVIDIA RTX 3090). Pre-compiling RNN computation graphs also decreased RNN inference time by 92.7% (average 91.4 ms; 95% CI, 79.4–105 ms). Third, lightweight language model decoding can further reduce latency. An existing phoneme-to-word software that uses efficient lookup-based data structures, along with unquantized LLM rescoring, reduced language model decoding time by 47.1% (average 469 ms; 95% CI, 455–484 ms).

### Deep ensembles improve offline decoding accuracy in various participants and performance regimes

After demonstrating the viability of deep ensemble-based decoding in real-time settings, we then evaluated whether its accuracy gains may translate to various participants, array configurations, and performance regimes. We used a diverse set of offline copy-speaking datasets from three participants (T12, T15, and T16) with microelectrode arrays in speech-related regions of the precentral gyrus (Extended Fig. 1; see Table 3 for a list of data collection sessions). For T12, the right hemisphere arrays were placed in a second surgery 1,155 days after the left hemisphere arrays. Since decoding accuracy substantially changed across these two timepoints (due to loss of spiking activity in the left hemisphere arrays over time), the recordings from each timepoint were evaluated separately (presented as “T12 left” and “T12 left + right”).

**Table 3.**
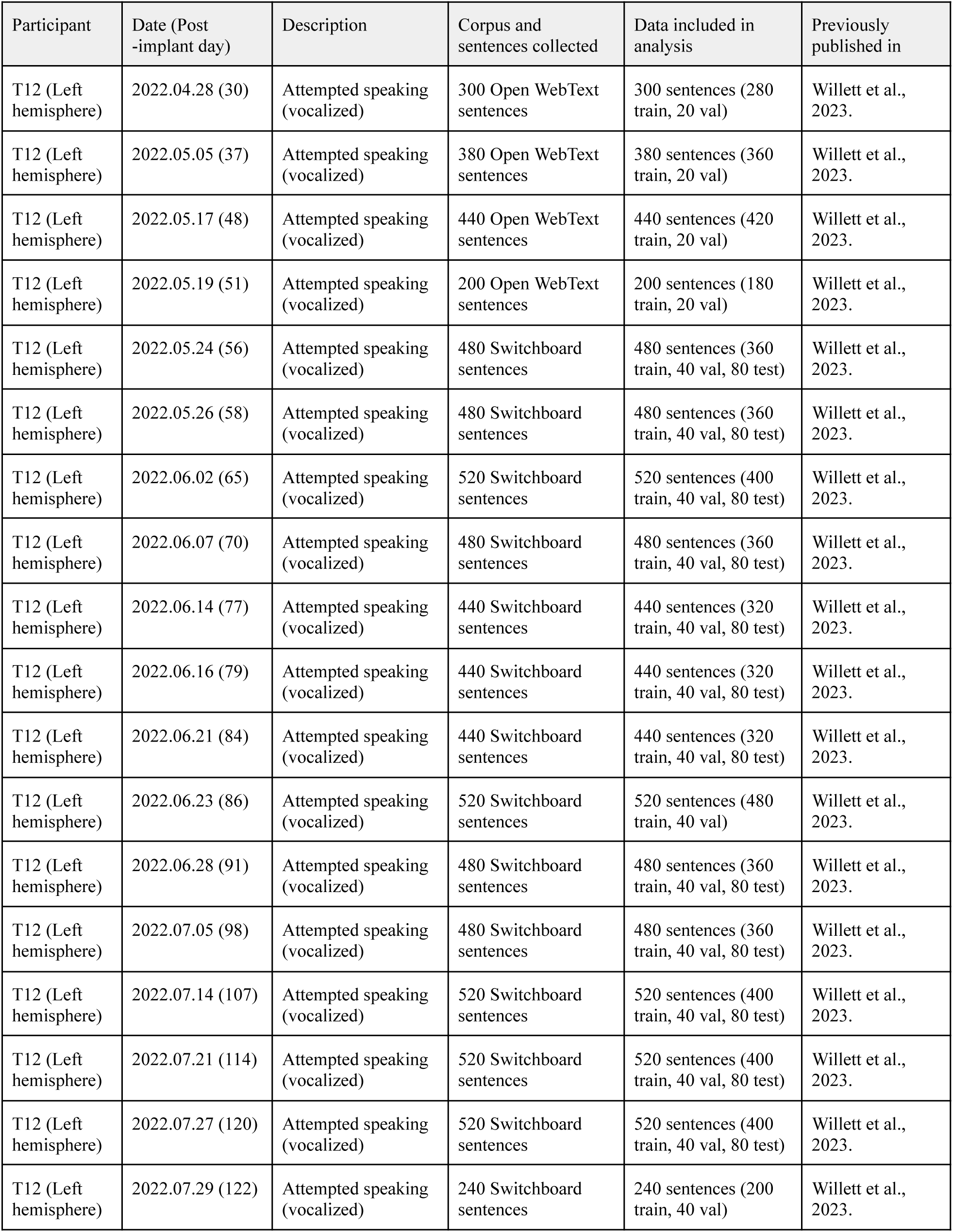
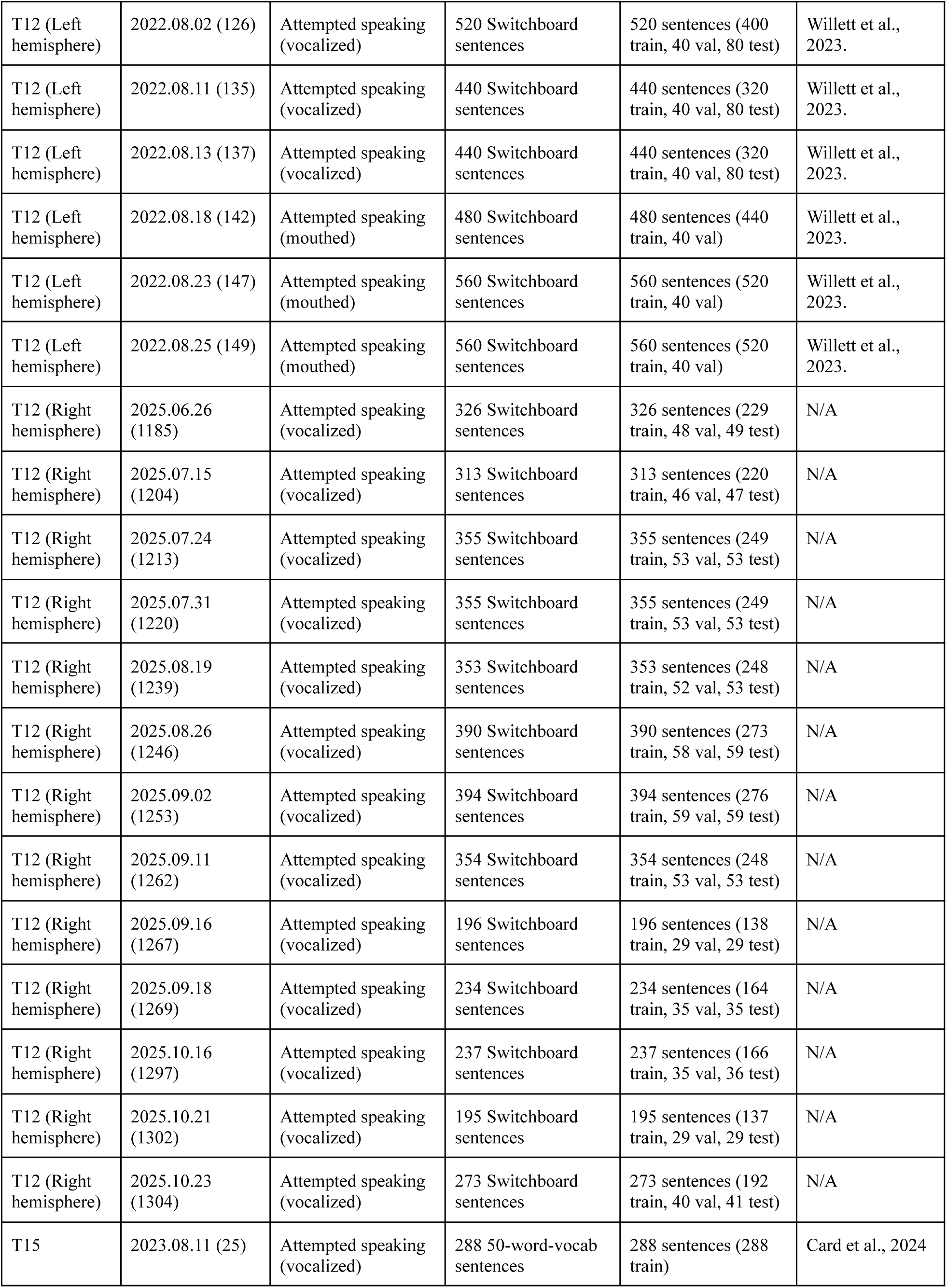
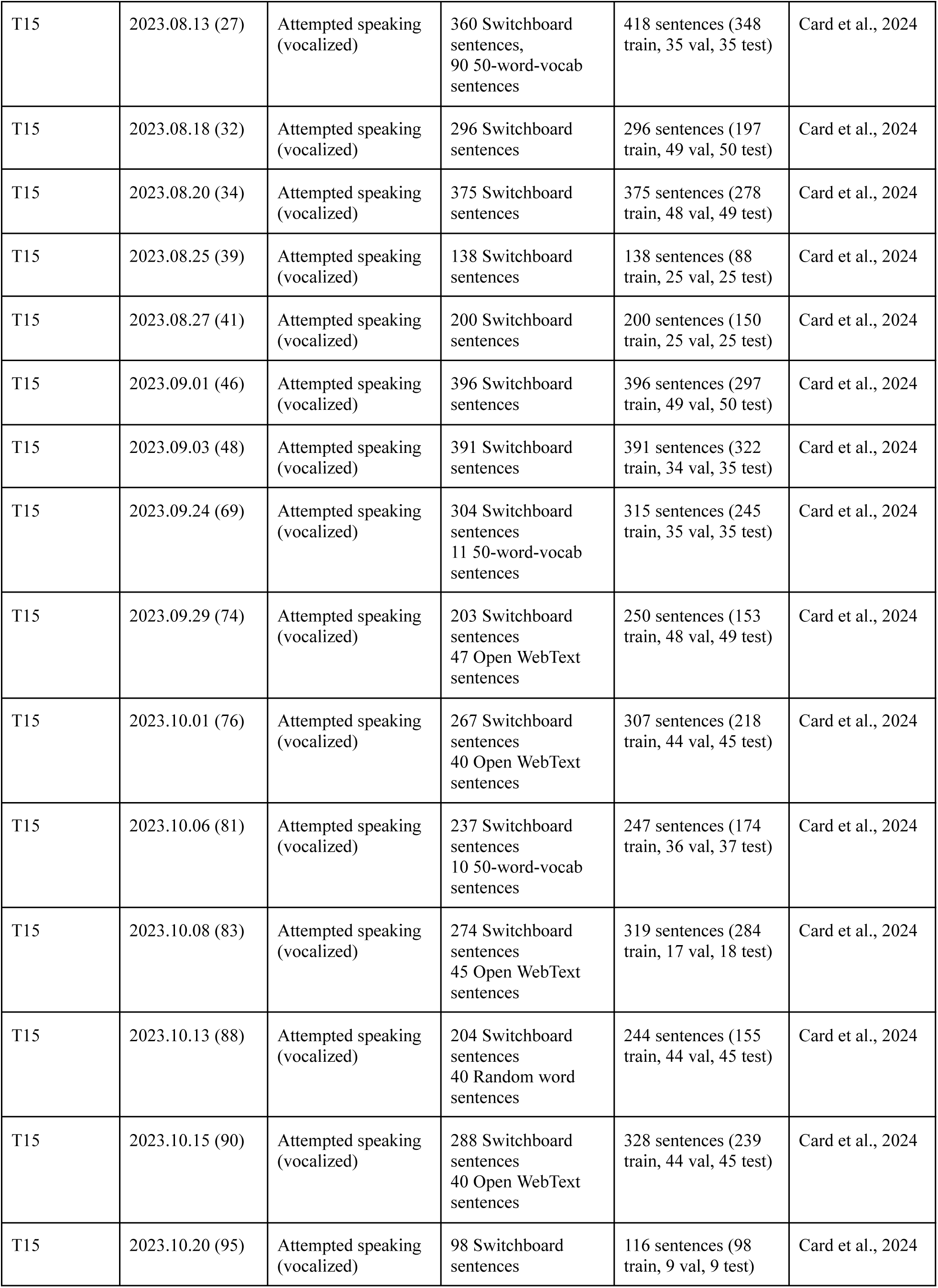
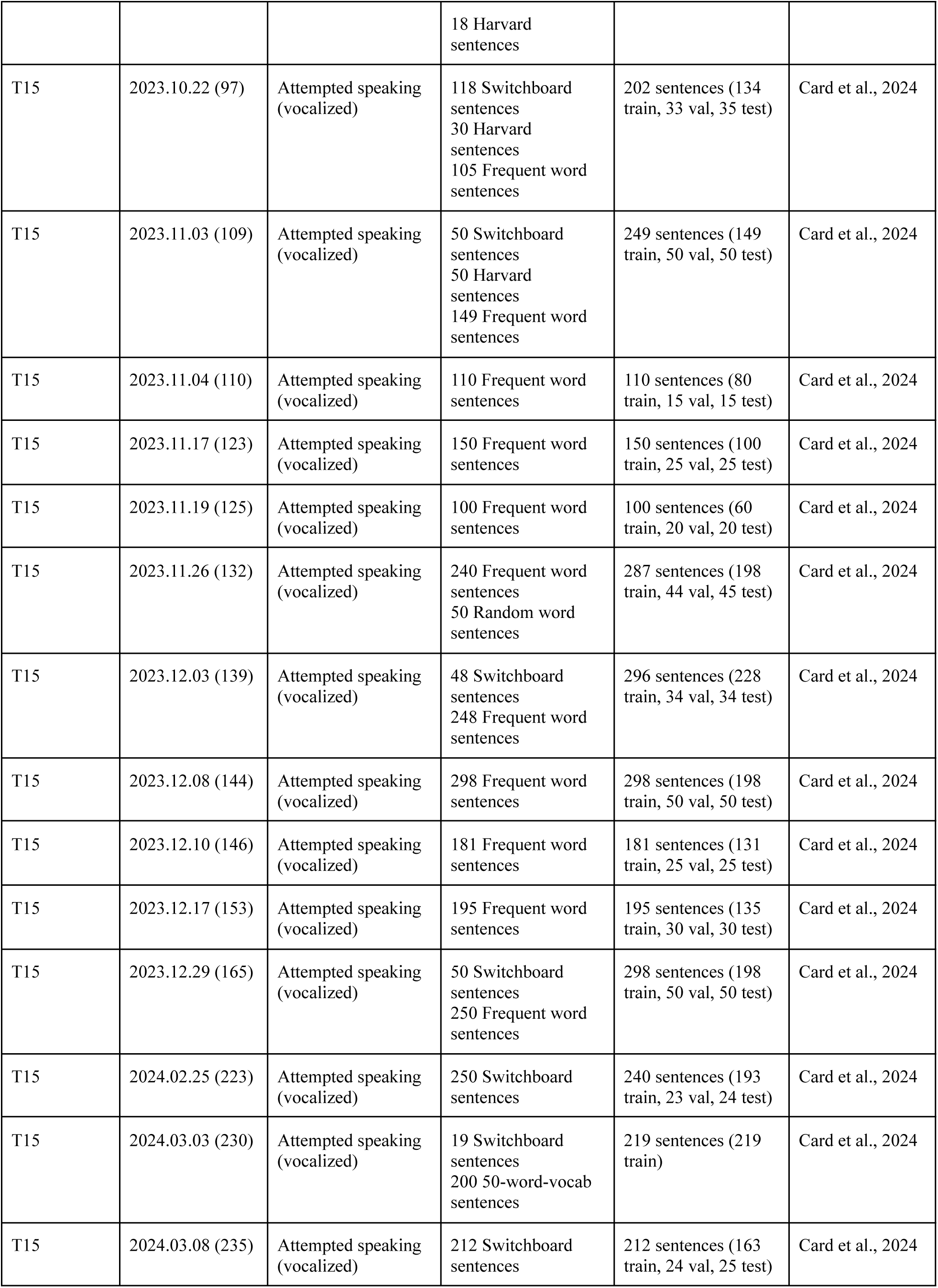
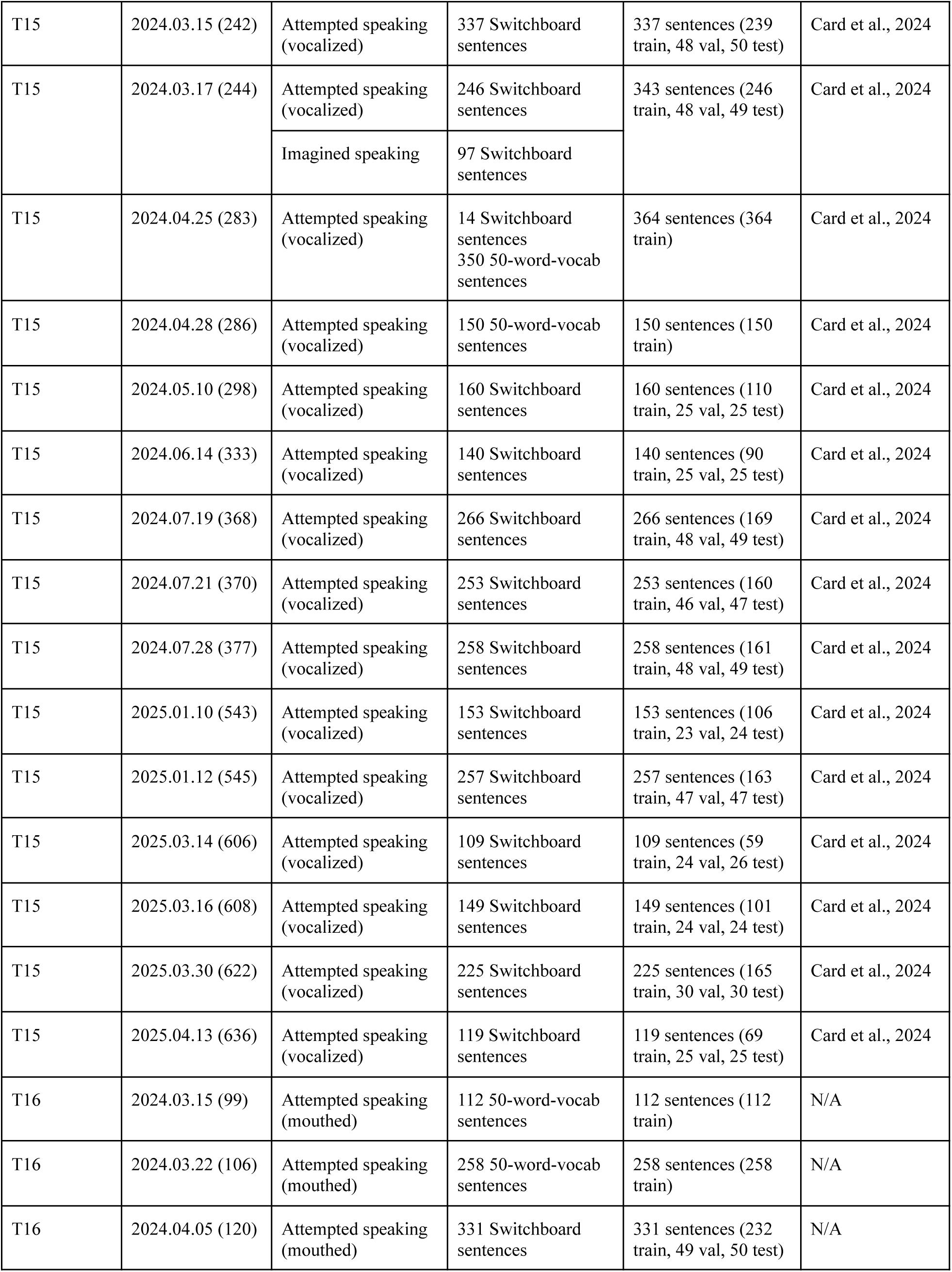
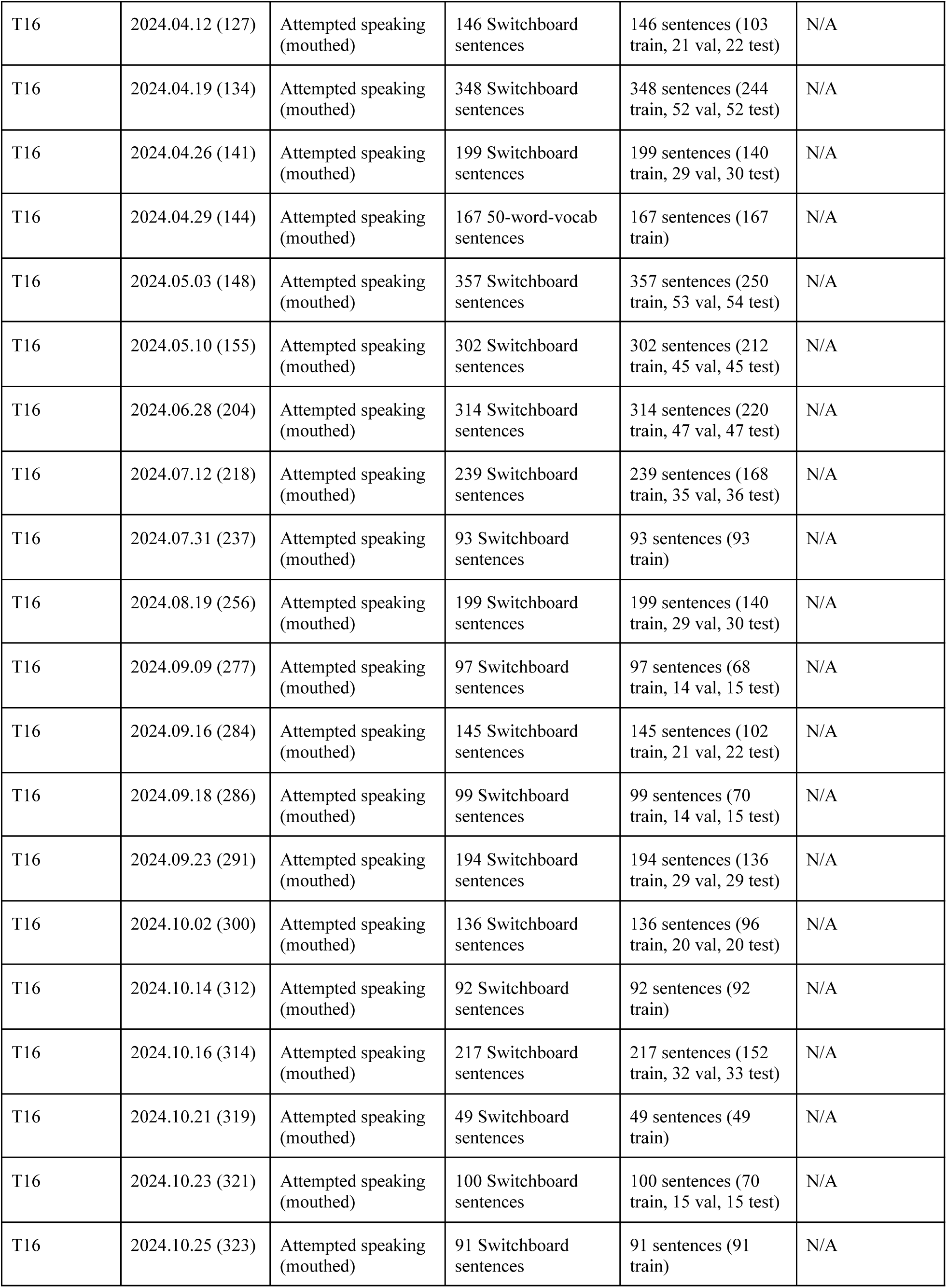
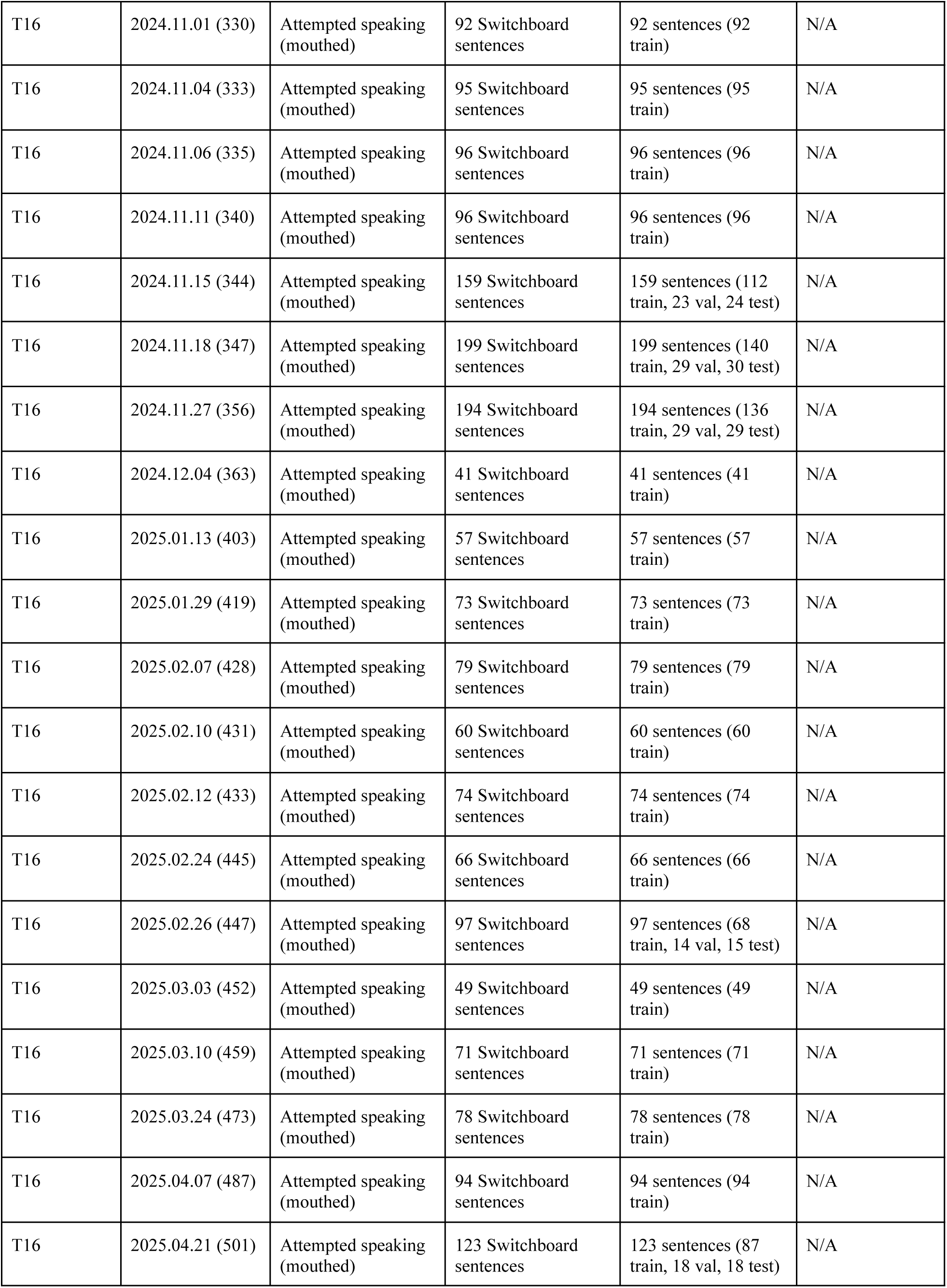
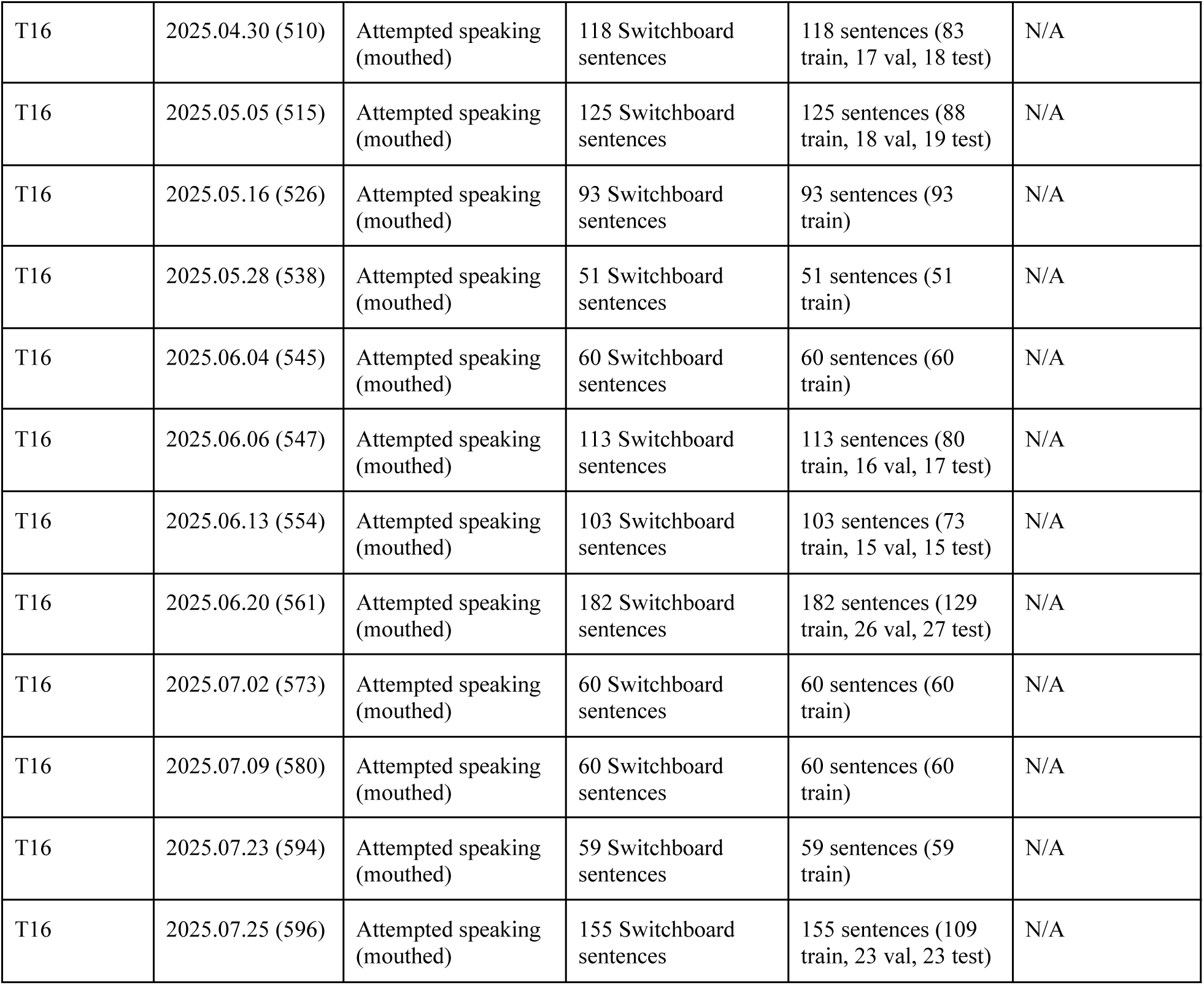
List of all data collection sessions used in this study.

**Table 4.** Decoder parameters for online decoding.

| Category | Name | Description | Value |
| --- | --- | --- | --- |
| Recurrent neural network offline training | # Units | Number of units in each GRU layer | 1280 |
|  | # Layers | Number of GRU layers | 5 |
|  | Kernel size | Number of input feature time bins stacked together as a single input for the RNN | 14 |
|  | Kernel stride | Number of time bins the RNN skips forward every inference step | 4 |
|  | L2 | L2 regularization factor | 1.0e−5 |
|  | Dropout | Dropout probability for GRU layers | 0.3 |
|  | Input # units | Number of units in each input layer | 768 |
|  | Input activation | Activation function of the input layer | softsign |
|  | Input dropout | Dropout probability for input layer | 0.25 |
|  | White noise SD | Standard deviation of white noise added to input data for regularization | 1.0 |
|  | Smooth kernel SD | Standard deviation of a temporal Gaussian smoothing kernel applied to neural data | 1.5 |
|  | Constant offset SD | Standard deviation of constant offset noise added to input data during training | 0.2 |
|  | Batch size | Number of sentences (trials) in each batch | 128 |
|  | Training batches | Number of training batches | 120000 |
|  | Patience | Number of batches to wait before halting training when either validation loss or phoneme error rate has not improved | 1000 |
|  | Validation batches | Number of training steps between validation runs | 100 |
|  | Learning rate start | Learning rate at the start of training | 1.0e-4 |
|  | Learning rate end | Target learning rate at the end of training | 0 |
|  | Learning rate decay | Type of learning rate decay | linear |
| | $\beta_1$ | ADAM gradient descent parameter (Exponential decay rate for first-moment estimate) | 0.9 |
| | $\beta_2$ | ADAM gradient descent parameter (Exponential decay rate for second-moment estimate) | 0.999 |
| | $\epsilon$ | ADAM gradient descent parameter (Constant added to denominator for numerical stability) | 1.0e-8 |
| Recurrent neural network online training | Learning rate start | Learning rate at the start of training | 2.0e-5 |
|  | Learning rate end | Target learning rate at the end of training | 1.0e-6 |
|  | Batch size | Number of sentences (trials) in each batch | 64 |
|  | New data percent | Percent of sentences (trials) per batch from new data from the current session | 0.7 |
|  | Min training steps | Minimum number of training steps to execute | 32 |
|  | Max training steps | Maximum number of training steps to execute | 200 |
|  | White noise SD | Standard deviation of white noise added to input data for regularization | 1.0 |
|  | Smooth kernel SD | Standard deviation of a temporal Gaussian smoothing kernel applied to neural data | 1.5 |
|  | Constant offset SD | Standard deviation of constant offset noise added to input data during training | 0.2 |
|  | Learning rate decay | Type of learning rate decay | linear |
| | $\beta_1$ | Adam gradient descent parameter (Exponential decay rate for first-moment estimate) | 0.9 |
| | $\beta_2$ | Adam gradient descent parameter (Exponential decay rate for second-moment estimate) | 0.999 |
| | $\epsilon$ | Adam gradient descent parameter (Constant added to denominator for numerical stability) | 1.0e-8 |
| 5-gram language model | Blank penalty | “Penalty” factor that the blank label probabilities are divided by | $\log(7)$ |
|  | Acoustic scale | Scaling factor on RNN’s log probabilities | 0.5 |
|  | Min active | Minimum active states during beam search | 200 |
|  | Max active | Maximum active states during beam search | 7000 |
|  | Beam | Beam size | 18 |
|  | n-best | Number of most likely sequences to return for LLM rescoring | 100 |
| Rescoring LLM | Acoustic scale | Scaling factor on RNN’s log probabilities | 0.5 |
|  | alpha | Interpolation weight between language models | 0.5 |

For each participant, we trained ten base decoders from random initializations and fine-tuned an open-source LLM (Qwen3-32B^44^) to aggregate ensemble predictions (Fig. 4a; see Tables 5 and 6 for a list of base decoder and merger LLM hyperparameters, respectively). Across all four datasets, deep ensembles significantly improved accuracy (Fig. 4c–f; 95% CIs of baseline and ensemble were non-overlapping) with relative WER reductions of 23.1% (T12 left hemisphere), 35.0% (T15), 31.1% (T16), and 27.6% (T12 left and right hemispheres).

**Table 5.** Base decoder parameters for offline analysis.

| T12 left hemisphere (Brain-to-Text ‘24) |  |  |  |
| --- | --- | --- | --- |
| Category | Name | Description | Value |
| Recurrent neural network training | # Units | Number of units in each GRU layer | 1024 |
|  | # Layers | Number of GRU layers | 5 |
|  | Kernel size | Number of input feature time bins stacked together as a single input for the RNN | 32 |
|  | Kernel stride | Number of time bins the RNN skips forward every inference step | 4 |
|  | Expanded prediction head | Whether the RNN used a deeper prediction head used introduced by the Linderman lab’s entry<sup>26</sup> | True |
|  | Bidirectional | Bidirectional GRU | True |
|  | GRU L2 | L2 regularization factor for GRU | 1.0e-5 |
|  | Input L2 | L2 regularization factor for input layer | 1.0e-5 |
|  | Dropout | Dropout probability for GRU layers | 0.4 |
|  | Input # units | Number of units in each input layer | 256 |
|  | Input activation | Activation function of the input layer | softsign |
|  | Input dropout | Dropout probability for input layer | 0.3 |
|  | Output dropout | Dropout probability for prediction layer | 0.4 |
|  | White noise SD | Standard deviation of white noise added to input data for regularization | 0.8 |
|  | Smooth kernel SD | Standard deviation of a temporal Gaussian smoothing kernel applied to neural data | 2.0 |
|  | Smooth kernel size | Number of time bins that the temporal Gaussian smoothing kernel is applied to | 20 |
|  | Constant offset SD | Standard deviation of constant offset noise added to input data during training | 0.2 |
|  | Batch size | Number of sentences (trials) in each batch | 64 |
|  | Training batches | Number of training batches | 20000 |
|  | Validation batches | Number of training steps between validation runs | 100 |
|  | Learning rate start | Learning rate at the start of training | 0.025 |
|  | Learning rate end | Target learning rate at the end of training | 0.0005 |
|  | Learning rate decay | Type of learning rate decay | linear |
|  | Optimizer type | Type of optimizer | Adam |
|  | CTC reduction type | How per-example CTC losses were aggregated across a batch. Sum returns sum of per-example losses; mean divides the CTC loss by the target sequence length | mean |
| | $\beta_1$ | Adam gradient descent parameter (Exponential decay rate for first-moment estimate) | 0.9 |
| | $\beta_2$ | Adam gradient descent parameter (Exponential decay rate for second-moment estimate) | 0.999 |
| | $\epsilon$ | Adam gradient descent parameter (Constant added to denominator for numerical stability) | 0.1 |
| 5-gram language model | Blank penalty | “Penalty” factor that the blank label probabilities are divided by | $\log(7)$ |
|  | Acoustic scale | Scaling factor on RNN’s log probabilities | 0.5 |
|  | Min active | Minimum active states during beam search | 200 |
|  | Max active | Maximum active states during beam search | 7000 |
|  | Beam | Beam size | 18 |
|  | n-best | Number of most likely sequences to return for LLM rescoring | 100 |
| Rescoring LLM | Acoustic scale | Scaling factor on RNN’s log probabilities | 0.5 |
|  | alpha | Interpolation weight between language models | 0.5 |

| T15 (Brain-to-Text ‘25) |  |  |  |
| --- | --- | --- | --- |
| Category | Name | Description | Value |
| Recurrent neural network training | # Units | Number of units in each GRU layer | 768 |
|  | # Layers | Number of GRU layers | 5 |
|  | Kernel size | Number of input feature time bins stacked together as a single input for the RNN | 32 |
|  | Kernel stride | Number of time bins the RNN skips forward every inference step | 4 |
|  | Expanded prediction head | Whether the RNN used a deeper prediction head used introduced by the Linderman lab’s entry<sup>26</sup> | True |
|  | Bidirectional | Bidirectional GRU | True |
| | GRU L2 | L2 regularization factor for GRU | $1.0\text{e-}5$ |
|  | Input L2 | L2 regularization factor for input layer | 0.1 |
|  | Dropout | Dropout probability for GRU layers | 0.14 |
|  | Input # units | Number of units in each input layer | 512 |
|  | Input activation | Activation function of the input layer | softsign |
|  | Input dropout | Dropout probability for input layer | 0.25 |
|  | Output dropout | Dropout probability for prediction layer | 0.22 |
|  | White noise SD | Standard deviation of white noise added to input data for regularization | 1.0 |
|  | Smooth kernel SD | Standard deviation of a temporal Gaussian smoothing kernel applied to neural data | 1.89 |
|  | Smooth kernel size | Number of time bins that the temporal Gaussian smoothing kernel is applied to | 100 |
|  | Constant offset SD | Standard deviation of constant offset noise added to input data during training | 0.2 |
|  | Batch size | Number of sentences (trials) in each batch | 64 |
|  | Training batches | Number of training batches | 60000 |
|  | Validation batches | Number of training steps between validation runs | 1000 |
|  | Learning rate start | Learning rate at the start of training | 5.0e−5 |
|  | Learning rate end | Target learning rate at the end of training | 1.0e−5 |
|  | Learning rate decay | Type of learning rate decay | cosine |
|  | Learning rate warmup batches | Number of training steps for learning rate warmup | 10000 |
|  | Optimizer type | Type of optimizer | AdamW |
|  | CTC reduction type | How per-example CTC losses were aggregated across a batch. Sum returns sum of per-example losses; mean divides the CTC loss by the target sequence length | sum |
| | $\beta_1$ | Adam gradient descent parameter (Exponential decay rate for first-moment estimate) | 0.9 |
| | $\beta_2$ | Adam gradient descent parameter (Exponential decay rate for second-moment estimate) | 0.999 |
| | $\epsilon$ | Adam gradient descent parameter (Constant added to denominator for numerical stability) | 1.0e−8 |
| 5-gram language model | Blank penalty | “Penalty” factor that the blank label probabilities are divided by | log(90) |
|  | Acoustic scale | Scaling factor on RNN’s log probabilities | 0.325 |
|  | Min active | Minimum active states during beam search | 200 |
|  | Max active | Maximum active states during beam search | 7000 |
|  | Beam | Beam size | 18 |
|  | n-best | Number of most likely sequences to return for LLM rescoring | 100 |
| Rescoring LLM | Acoustic scale | Scaling factor on RNN's log probabilities | 0.235 |
|  | alpha | Interpolation weight between language models | 0.5 |

| T16 |  |  |  |
| --- | --- | --- | --- |
| Category | Name | Description | Value |
| Recurrent neural network training | # Units | Number of units in each GRU layer | 1024 |
|  | # Layers | Number of GRU layers | 5 |
|  | Kernel size | Number of input feature time bins stacked together as a single input for the RNN | 32 |
|  | Kernel stride | Number of time bins the RNN skips forward every inference step | 4 |
|  | Expanded prediction head | Whether the RNN used a deeper prediction head used introduced by the Linderman lab's entry<sup>26</sup> | True |
|  | Bidirectional | Bidirectional GRU | True |
|  | GRU L2 | L2 regularization factor for GRU | 1.0e-5 |
|  | Input L2 | L2 regularization factor for input layer | 1.0e-5 |
|  | Dropout | Dropout probability for GRU layers | 0.4 |
|  | Input # units | Number of units in each input layer | 128 |
|  | Input activation | Activation function of the input layer | softsign |
|  | Input dropout | Dropout probability for input layer | 0.3 |
|  | Output dropout | Dropout probability for prediction layer | 0.4 |
|  | White noise SD | Standard deviation of white noise added to input data for regularization | 0.8 |
|  | Smooth kernel SD | Standard deviation of a temporal Gaussian smoothing kernel applied to neural data | 2.0 |
|  | Smooth kernel size | Number of time bins that the temporal Gaussian smoothing kernel is applied to | 40 |
|  | Constant offset SD | Standard deviation of constant offset noise added to input data during training | 0.2 |
|  | Batch size | Number of sentences (trials) in each batch | 64 |
|  | Training batches | Number of training batches | 20000 |
|  | Validation batches | Number of training steps between validation runs | 100 |
|  | Learning rate start | Learning rate at the start of training | 2.0e−4 |
|  | Learning rate end | Target learning rate at the end of training | 5.0e−5 |
|  | Day learning rate start | Learning rate for the input layer at the start of training | 1.0e−4 |
|  | Day learning rate end | Target learning rate for the input layer at the end of training | 5.0e−5 |
|  | Learning rate decay | Type of learning rate decay | cosine |
|  | Learning rate warmup batches | Number of training steps for learning rate warmup | 2000 |
|  | Optimizer type | Type of optimizer | AdamW |
|  | CTC reduction type | How per-example CTC losses were aggregated across a batch. Sum returns sum of per-example losses; mean divides the CTC loss by the target sequence length | mean |
| | $\beta_1$ | Adam gradient descent parameter (Exponential decay rate for first-moment estimate) | 0.9 |
| | $\beta_2$ | Adam gradient descent parameter (Exponential decay rate for second-moment estimate) | 0.999 |
| | $\epsilon$ | Adam gradient descent parameter (Constant added to denominator for numerical stability) | 1.0e−8 |
| 5-gram language model | Blank penalty | “Penalty” factor that the blank label probabilities are divided by | log(6) |
|  | Acoustic scale | Scaling factor on RNN’s log probabilities | 0.25 |
|  | Min active | Minimum active states during beam search | 200 |
|  | Max active | Maximum active states during beam search | 7000 |
|  | Beam | Beam size | 18 |
|  | n-best | Number of most likely sequences to return for LLM rescoring | 100 |
| Rescoring LLM | Acoustic scale | Scaling factor on RNN’s log probabilities | 0.25 |
|  | alpha | Interpolation weight between language models | 0.7 |

| T12 left right hemispheres |  |  |  |
| --- | --- | --- | --- |
| Category | Name | Description | Value |
| Recurrent neural network training | # Units | Number of units in each GRU layer | 1280 |
|  | # Layers | Number of GRU layers | 5 |
|  | Kernel size | Number of input feature time bins stacked together as a single input for the RNN | 32 |
|  | Kernel stride | Number of time bins the RNN skips forward every inference step | 4 |
|  | Expanded prediction head | Whether the RNN used a deeper prediction head used introduced by the Linderman lab's entry<sup>26</sup> | False |
|  | Bidirectional | Bidirectional GRU | True |
|  | GRU L2 | L2 regularization factor for GRU | 0.1 |
|  | Input L2 | L2 regularization factor for input layer | 0.001 |
|  | Dropout | Dropout probability for GRU layers | 0.4 |
|  | Input # units | Number of units in each input layer | 768 |
|  | Input activation | Activation function of the input layer | softsign |
|  | Input dropout | Dropout probability for input layer | 0.4 |
|  | Output dropout | Dropout probability for prediction layer | 0.2 |
|  | White noise SD | Standard deviation of white noise added to input data for regularization | 0.9 |
|  | Smooth kernel SD | Standard deviation of a temporal Gaussian smoothing kernel applied to neural data | 1.75 |
|  | Smooth kernel size | Number of time bins that the temporal Gaussian smoothing kernel is applied to | 40 |
|  | Constant offset SD | Standard deviation of constant offset noise added to input data during training | 0.15 |
|  | Batch size | Number of sentences (trials) in each batch | 64 |
|  | Training batches | Number of training batches | 120000 |
|  | Validation batches | Number of training steps between validation runs | 500 |
|  | Learning rate start | Learning rate at the start of training | 5.0e-5 |
|  | Learning rate end | Target learning rate at the end of training | 1.0e-5 |
|  | Learning rate decay | Type of learning rate decay | cosine |
|  | Learning rate warmup batches | Number of training steps for learning rate warmup | 10000 |
|  | Optimizer type | Type of optimizer | AdamW |
|  | CTC reduction type | How per-example CTC losses were aggregated across a batch. Sum returns sum of per-example losses; mean divides the CTC loss by the target sequence length | sum |
| | $\beta_1$ | Adam gradient descent parameter (Exponential decay rate for first-moment estimate) | 0.9 |
| | $\beta_2$ | Adam gradient descent parameter (Exponential decay rate for second-moment estimate) | 0.999 |
| | $\epsilon$ | Adam gradient descent parameter (Constant added to denominator for numerical stability) | 1.0e-8 |
| Recurrent neural network fine-tuning on right hemisphere data<br>(If hyperparameters not shown here, we used the same values as the RNN training table above) | GRU L2 | L2 regularization factor for GRU | 0.1 |
|  | Input L2 | L2 regularization factor for input layer | 0.001 |
|  | Dropout | Dropout probability for GRU layers | 0.4 |
|  | Input activation | Activation function of the input layer | softsign |
|  | Input dropout | Dropout probability for input layer | 0.4 |
|  | Output dropout | Dropout probability for prediction layer | 0.22 |
|  | White noise SD | Standard deviation of white noise added to input data for regularization | 1.0 |
|  | Smooth kernel SD | Standard deviation of a temporal Gaussian smoothing kernel applied to neural data | 1.89 |
|  | Smooth kernel size | Number of time bins that the temporal Gaussian smoothing kernel is applied to | 100 |
|  | Constant offset SD | Standard deviation of constant offset noise added to input data during training | 0.2 |
|  | Batch size | Number of sentences (trials) in each batch | 64 |
|  | Training batches | Number of training batches | 10000 |
|  | Validation batches | Number of training steps between validation runs | 100 |
|  | Learning rate start | Learning rate at the start of training | 8.0e-06 |
|  | Learning rate end | Target learning rate at the end of training | 1.0e-06 |
|  | Learning rate decay | Type of learning rate decay | linear |
| 5-gram language model | Blank penalty | “Penalty” factor that the blank label probabilities are divided by | log(7) |
|  | Acoustic scale | Scaling factor on RNN’s log probabilities | 0.5 |
|  | Min active | Minimum active states during beam search | 200 |
|  | Max active | Maximum active states during beam search | 7000 |
|  | Beam | Beam size | 18 |
|  | n-best | Number of most likely sequences to return for LLM rescoring | 100 |
| Rescoring LLM | Acoustic scale | Scaling factor on RNN's log probabilities | 0.5 |
|  | alpha | Interpolation weight between language models | 0.5 |

**Table 6.**
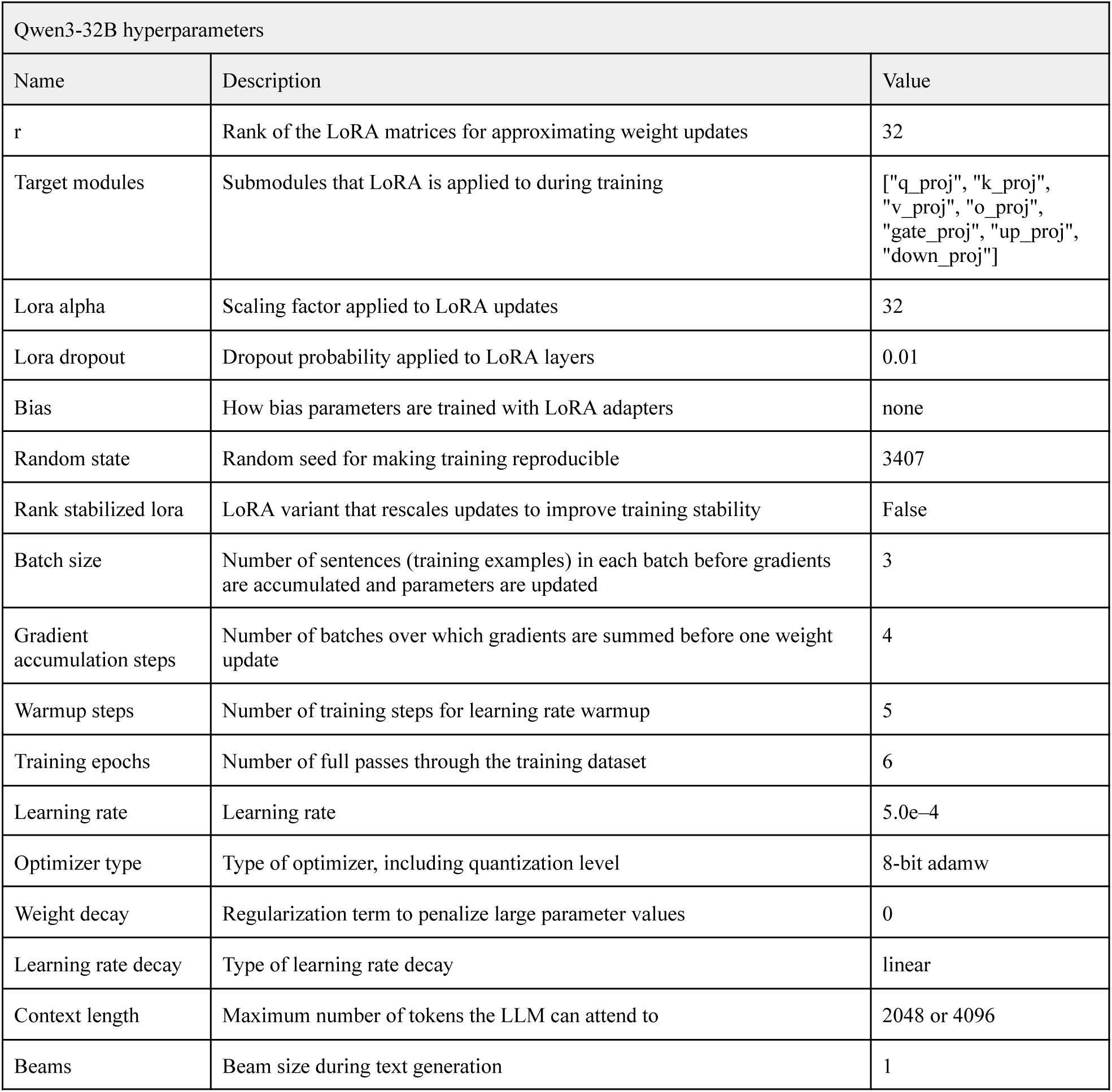
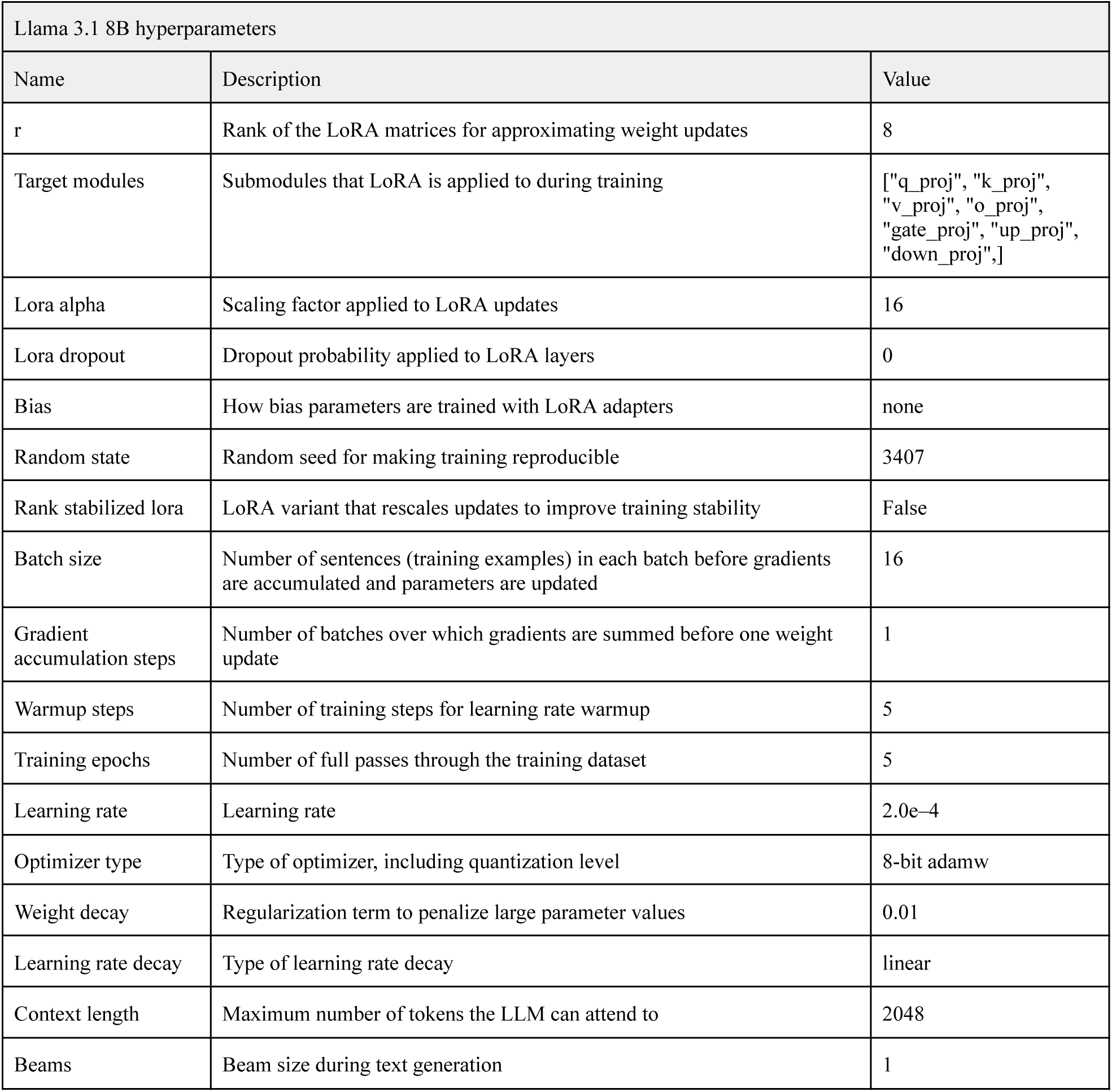
Merger LLM hyperparameters for offline analysis.

**Table 7.** Merger LLM hyperparameter search space.

| Name | Description | Value |
| --- | --- | --- |
| r | Rank of the LoRA matrices for approximating weight updates | 8, 16, 32, 64 |
| Lora alpha/r | Multiplicative factor used to set the LoRA update scaling parameter (alpha) relative to the LoRA matrix rank (r). For example, a r of 16 and alpha/r of 2 yielded an alpha factor of $16 \times 2 = 32$ | 1, 2, 4 |
| Lora dropout | Dropout probability applied to LoRA layers | 0.01, 0.02, 0.05 |
| Bias | How bias parameters are trained with LoRA adapters | all, none, lora_only |
| Batch size | Number of sentences (training examples) in each batch before gradients are accumulated and parameters are updated | 1, 2, 3, 4 |
| Gradient accumulation steps | Number of batches over which gradients are summed before one weight update | 1, 2, 4, 8 |
| Warmup steps | Number of training steps for learning rate warmup | 0, 5, 10, 50 |
| Training epochs | Number of full passes through the training dataset | 1, 3, 6 |
| Learning rate | Learning rate | 1e-03, 8e-04, 5e-04, 3e-04, 2e-04, 1e-04, 5e-05 |
| Weight decay | Regularization term to penalize large parameter values | 0, 0.01, 1e-03, 1e-05 |
| Learning rate decay | Type of learning rate decay | linear, cosine |

### Test-time augmentation serves as a lightweight alternative to ensembles in brain-to-text decoding

Although our cloud framework offers a scalable way for deploying ensemble speech BCIs, it may still be challenging in more resource-constrained scenarios. The main memory bottleneck is the need for continuous recalibration and inference of multiple base decoders. Therefore, we next developed a computationally efficient “pseudoensembling” method using test-time augmentation, a technique that approximates ensemble behavior by using a single base decoder to generate diverse predictions from multiple perturbed versions of the same input (Fig. 4b). This method is commonly used in computer vision^45–47^ to improve accuracy without ensembling overhead. Specifically, we injected randomly initialized Gaussian white noise into neural data, and predictions generated by each perturbed version were subsequently merged using a fine-tuned LLM^42,44,48^. For each session day, noise strength was selected by sweeping across noise levels to find the noise level that generated a diversity of outputs that best matched the diversity of multiple independently trained base decoders (diversity was defined as word-level disagreement frequency on a separate validation set^49^). Across all four datasets, our pseudoensembling method consistently improved accuracy (Fig. 4c–f; 95% CIs of baseline and pseudoensemble were non-overlapping) with relative error rate reductions of 15.7% (T12 left hemisphere), 17.0% (T15), 11.7% (T16), and 9.4% (T12 left+right hemisphere), although full ensembles achieved larger performance gains.

### Design considerations for ensemble-based speech BCIs

Finally, we explored three design considerations for ensemble-based speech BCIs: hypothesis aggregation method, ensemble size, and training dataset size.

First, we compared aggregation methods including various families of fine-tuned LLMs^42,44,48,50,51^ and a non-LLM heuristic (ROVER^33^), which performs word position-wise voting after aligning candidate hypotheses into a word lattice. All aggregation methods consistently improved upon the baseline across all four participants (Fig. 5a–d). Surprisingly, the performance gains via heuristic-based aggregation approached those of smaller LLMs at times, offering a viable alternative when compute is constrained or minimizing latency is critical. However, more powerful LLMs yielded better hypothesis aggregation. Given the exponential improvements in intelligence per parameter size of open-source LLMs^52^, this suggests that further performance gains may be achievable with lower computational cost, although it is unlikely that more powerful LLMs can indefinitely improve ensemble accuracy. Furthermore, consistent with online results, ensembling across multiple fine-tuning checkpoints of the merger LLM improved performance, even when using a simple selection rule based on output probability.

**Fig. 5.**
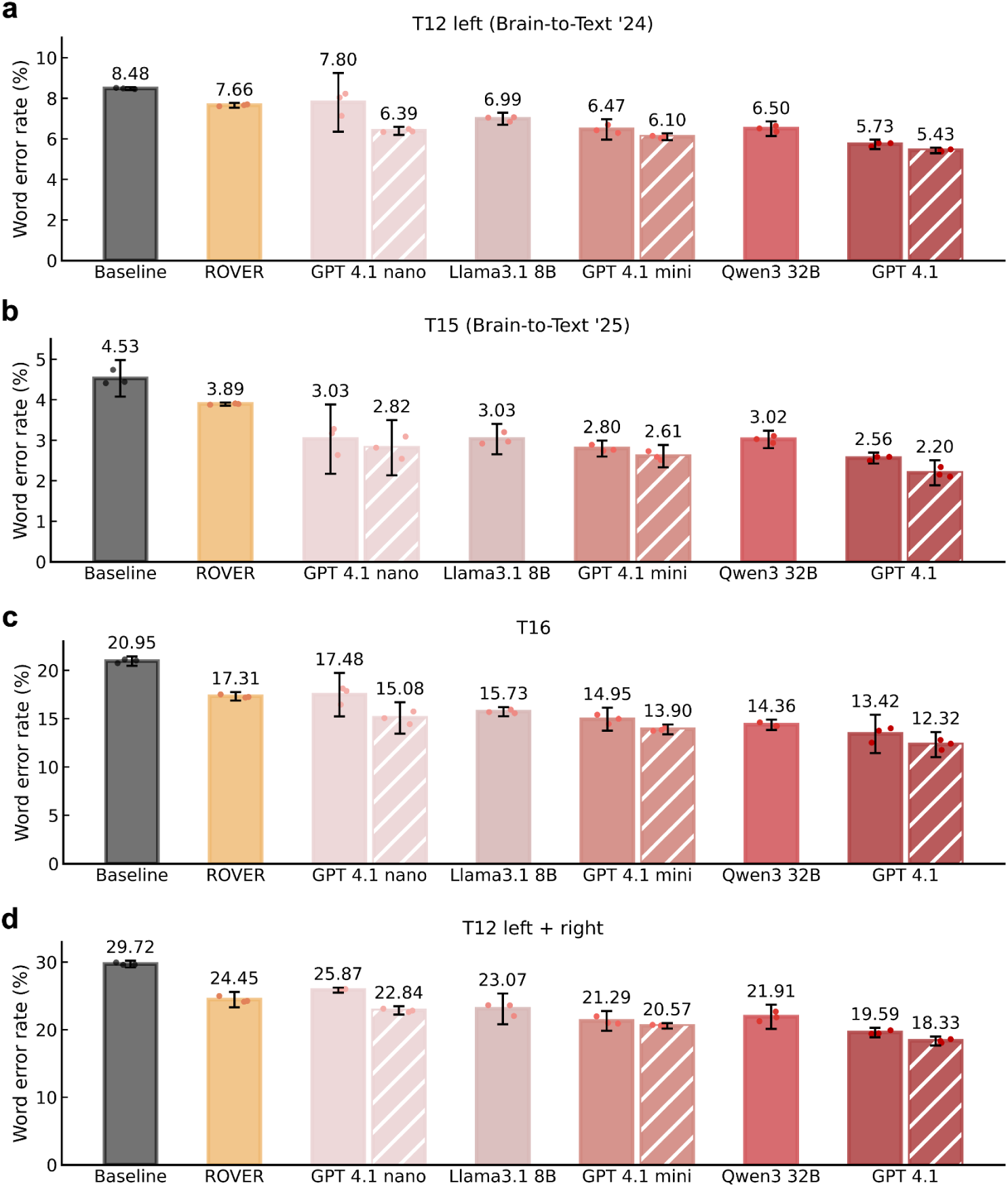
Decoding performance comparison of different hypothesis aggregation methods. **a,** Decoding performance of deep ensembles on T12 left hemisphere data using fine-tuned LLMs of varying sizes and a non-LLM heuristic (ROVER). Hatched bars denote checkpoint ensembling (selecting the output with the highest probability across predictions from multiple fine-tuning checkpoints of the merger LLM). Results are compared against the single-decoder baseline (gray bar). Bar heights indicate mean word error rate across three training initializations. Error bars represent 95% CIs. **b–d,** Same as in **a,** but for T15, T16, and T12 right hemisphere data, respectively.

Next, we investigated how ensemble performance scaled with the amount of training data used to fine-tune the merger LLM, as training data is often scarce in clinical translation settings^53^. Across all four participants, the deep ensemble outperformed the baseline even when the merger LLM was fine-tuned on as few as 20 sentences, and performance further improved as more training data was available (Fig. 6a–d). Furthermore, the fact that the merger LLM was able to learn the base decoder output distributions with minimal training data also opens the possibility that relatively rapid online recalibration of the merger LLM may be feasible, in the event of a sudden distribution shift.

**Fig. 6.**
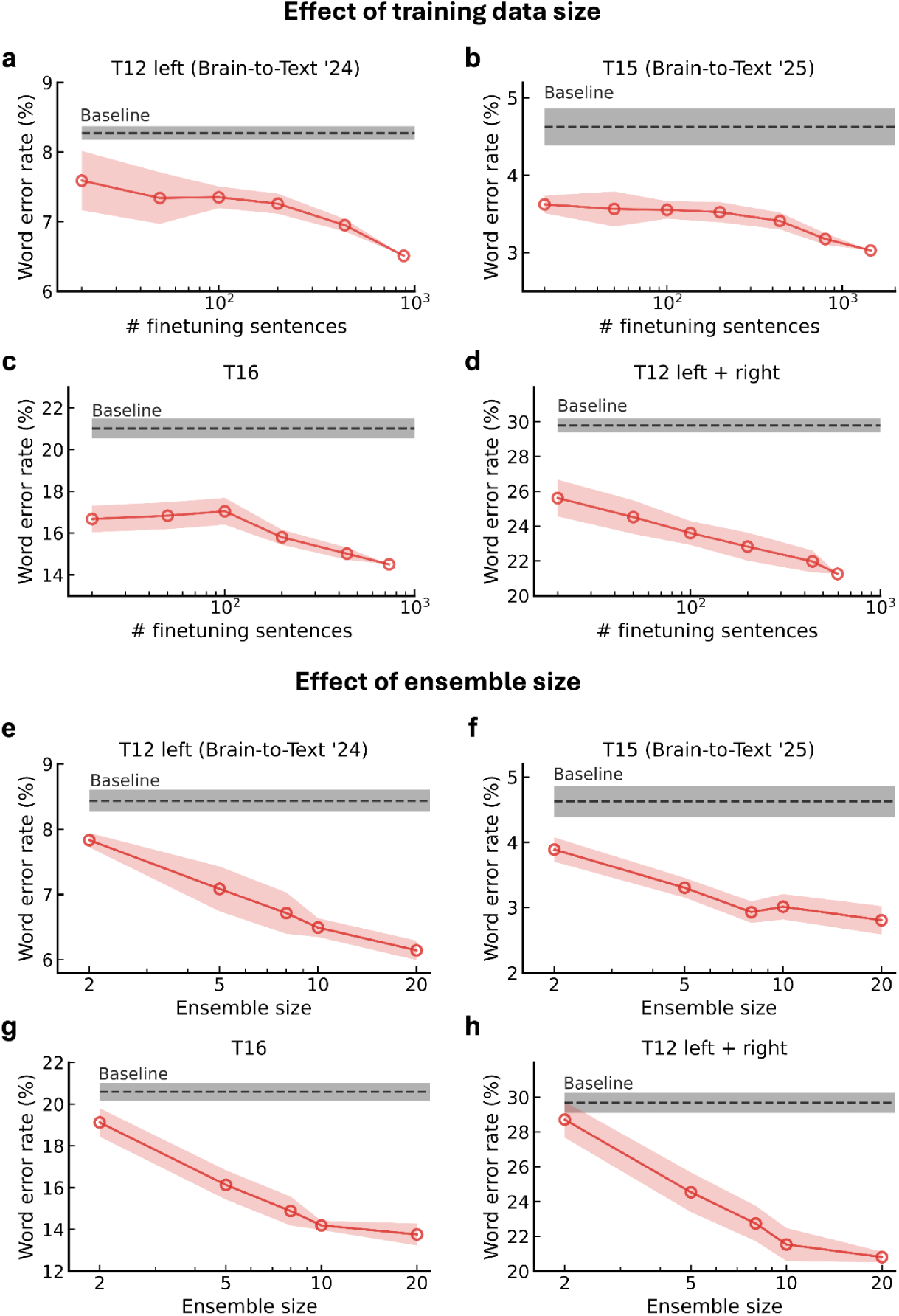
Scaling properties of ensemble-based speech BCIs. **a,** Word error rate as a function of the merger LLM fine-tuning dataset size for T12 left hemisphere data, obtained by randomly sampling a subset of the fine-tuning dataset and re-training the merger LLM. Each point represents performance averaged across ten independent subsamples. The black dashed line denotes the single-model baseline. The shaded regions represent 95% CIs. **b–d,** Same as in **a,** but for T15, T16, and T12 right hemisphere data, respectively. **e,** Word error rate as a function of ensemble size for T12 left hemisphere data, obtained by re-training a deep ensemble of given size. Each point represents performance averaged across five distinct random initializations during training. The black dashed line denotes the single-model baseline. The shaded regions represent 95% CIs. **f–h,** Same as in **e,** but for T15, T16, and T12 right hemisphere data, respectively.

Finally, we investigated how performance scaled with ensemble size, as performing parallel recalibration and inference of multiple base decoders was the primary source of test-time compute in our ensemble-based BCI. Performance improved even with as few as two base decoders and increased with more, following a roughly log-linear trend (Fig. 6e–h), although the gains are unlikely to indefinitely improve as additional decoders contribute increasingly redundant predictions. This suggests that ensemble sizes may be flexibly adjusted to deployment constraints as needed.

## Discussion

Deep ensembles have consistently shown substantial accuracy improvements in offline neural decoding competitions^24–31^, yet whether these results can translate to real-time use was previously untested. Here, we showed that the accuracy gains of deep ensembles translate effectively into real-time, closed-loop operation, reducing WER from 33.7% to 26.0% relative to the previous state-of-the-art method demonstrated in real-time^3,4,6,7^. Importantly, our method improved performance to a range considered sufficient for minimally functional communication^16^. Consistent with these real-time findings, we further found offline that deep ensembles improve performance across multiple participants, array configurations, baseline error regimes, amounts of training data, ensemble sizes, and hypothesis aggregation methods, suggesting that ensemble speech BCIs may be broadly useful across diverse deployment settings.

More broadly, our results suggest a consistent performance trend for speech BCIs: accuracy tends to improve as greater test-time compute is allocated, such as generating more diverse candidate predictions or combining them with a more capable merger model. We have further shown that these compute-driven gains can be realized in real-time using cloud resources, through the first closed-loop demonstration of a cloud-based BCI and practical implementation principles to significantly reduce inference latency. Overall, these results raise the possibility that future BCIs could be designed with minimal local resources, instead using scalable cloud infrastructure that can enable more powerful decoders such as deep ensembles. Looking ahead, as hardware and AI inference efficiency rapidly improves^52,54^, cloud-supported ensemble BCIs may further excel in performance and cost-effectiveness.

While cloud-based ensembles can enhance functionality, reliable network access or cloud resources may be limited in many clinical translation settings^55,56^. To address this issue, we also proposed a computationally efficient pseudoensembling method that can partially retain the performance of deep ensembles. Importantly, this method does not require the simultaneous recalibration and inference of multiple base decoders, which was the primary computation cost of our ensemble speech BCI. Since inference using this pseudoensembling approach can be easily parallelized, it could be implemented on the same local hardware setup provisioned for inference in prior real-time state-of-the-art systems^4–7^ (see Extended Fig. 2b for a potential system diagram). Together, these results offer a highly versatile framework that can improve decoding accuracy across a clinically relevant continuum of deployment scenarios.

We believe the broader implications of our results for communication BCIs are promising. In principle, because our ensemble speech BCI operates on word-level decoder outputs, it could be extended to a broader range of brain-to-text communication systems. For example, prior work has demonstrated BCIs decoding handwriting^57,58^, keyboard-based typing^59,60^, and keyword-based spelling^61^. These various communication BCIs that produce discrete text-based outputs could benefit from ensembling, especially as we have shown that deep ensembles can be implemented in real-time as a modular post-processing component, without modification of the base decoder, and using off-the-shelf libraries. Furthermore, the use of an LLM merger makes it easy to incorporate conversational context into hypothesis aggregation, which has preliminarily been shown to improve decoding accuracy^62^.

Deep ensembles are also fundamentally model-agnostic. Recent works in brain-to-text decoding have explored more hardware-efficient end-to-end decoding architectures^31,36^, recalibration methods^29,63^, and phoneme-to-word decoding^30,64^. While our cloud-based implementation can indeed support more computationally intensive ensembles, with continued improvements in decoding methods, alongside hardware developments, ensemble-based decoding may become increasingly practical under smaller compute budgets. Future work could explore how such architectural advances can be combined with deep ensembles or pseudoensembles to further improve accuracy and real-time deployability.

However, a limitation of this study is that real-time performance was evaluated for a single participant. Although our offline analysis extends our results to a broader error rate regime and array configurations, real-time validation across additional participants would help further confirm the generality of our findings. In addition, real-time performance was evaluated over a relatively limited time period. Work remains to determine the long-term stability of ensemble-based speech BCIs, including everyday-use settings, and how these systems could efficiently adapt to nonstationarities over time.

For individuals whose primary means of communication is a speech BCI, even modest improvements in decoding accuracy may translate into meaningful gains in communication reliability. By demonstrating practical strategies and considerations for designing ensemble-based decoders under real-time constraints, we believe this work will help to bring communication BCIs closer to widespread clinical adoption.

## Data and code availability

De-identified data and the code used in the analyses will be available upon publication.

## Acknowledgements

We thank participants T12, T15, and T16 and their care partners for their generous time and contributions to this research. We also appreciate the administrative support from B. Davis, K. Tsou, S. Kosasih, M. Masood, B. Travers, and D. Rosler for clinical site oversight.

This work was supported by an ALS Pilot Clinical Trial Award (AL220043) from the Office of the Assistant Secretary of Defense for Health Affairs; a New Innovator Award (NIH 1DP2DC021055) from the National Institutes of Health, managed by the National Institute on Deafness and Other Communication Disorders; a grant (872146SPI) from the Simons Collaboration for the Global Brain; a postdoctoral fellowship from the A.P. Giannini Foundation; support from the Office of Research and Development, Department of Veterans Affairs (nos. N2864C, A2295R, and A4820R); the Wu Tsai Neurosciences Institute; the Howard Hughes Medical Institute; Larry and Pamela Garlick; NIDCD (nos. U01DC017844, R01DC014034); Society for Neuroscience Trainee Professional Development Award; the Bruce and Elizabeth Dunlevie Stanford Bio-X Interdisciplinary Graduate Fellowship; NIH F32HD112173; the Searle Scholars Program; Career Awards at the Scientific Interface from the Burroughs Wellcome Fund; NIH-NINDS/OD DP2NS127291 and the Simons Foundation as part of the Simons-Emory International Consortium on Motor Control; and the NSF GRFP.

The content is solely the responsibility of the authors and does not necessarily represent the official views of the National Institutes of Health, or the Department of Veterans Affairs, or the United States Government. CAUTION: Investigational Device. Limited by Federal Law to Investigational Use.

## Author contributions

S.Y. and F.R.W. conceived the study and led the development of all experiments. S.Y. led the interpretation and analysis of all experiments, and wrote the manuscript with input from D.R.D.

S.Y., D.T.A., S.M., A.D.L., C.F., and B.A.K. developed the ensemble speech BCI software.

D.T.A. and S.Y. developed the data collection rig software at the Stanford University site.

A.S., C.V., and N.H. were responsible for data collection, session scheduling, logistics, and daily equipment setup and disconnection for participant T12, and A.D.L. assisted with preparation for data collection.

N.S.C. and Z.F. contributed to the offline analysis code.

N.S.C., M.W., S.R.N.-T., and B.G.J. developed and executed the experiments at their respective sites.

C.I., P.H.B., S.R.N.-T. were responsible for coordination of session scheduling, logistics and daily equipment setup/disconnection for participants T15 and T16 respectively.

L.R.H. is the sponsor-investigator of the multisite BrainGate2 pilot clinical trial.

J.M.H. planned and performed both T12’s left and right hemisphere array placement surgery and was responsible for all clinical trial-related activities at Stanford University.

D.M.B. planned and performed T15’s array placement surgery and was responsible for all clinical trial-related activities at University of California, Davis. S.D.S. and D.M.B. supervised and guided all research activities at UC Davis.

N.A.Y. planned and performed T16’s array placement surgery and was responsible for all clinical trial-related activities at Emory University. C.P. and N.A.Y. supervised and guided all research activities at Emory.

The study was supervised and guided by J.M.H. and F.R.W. All authors reviewed and edited the manuscript.

## Methods

### Neural recordings and data

#### Study participants and approval

This study includes three participants, referred to as T12, T15, and T16, all of whom gave informed consent and were enrolled in the BrainGate2 Neural Interface System pilot clinical trial (ClinicalTrials.gov Identifier: NCT00912041, registered June 3, 2009). Approval for this pilot clinical trial was granted under an Investigational Device Exemption (IDE) by the US Food and Drug Administration (Investigational Device Exemption #G090003), as well as the Institutional Review Boards of Stanford University (protocol #52060), the University of California Davis (protocol #1843264), and Emory University (protocol #STUDY00003070). All research was performed in accordance with relevant guidelines and regulations.

T12, a left-handed woman with slowly-progressive bulbar-onset Amyotrophic Lateral Sclerosis (ALS), was 68–71 years old at the time of data collection. She was diagnosed at age 59 (ALS-FRS score of 26 at the time of study enrollment). She had eight 64-channel microelectrode arrays placed in speech-related cortex of both hemispheres. In the left hemisphere, she had two arrays in HCP-identified^65^ area 6v of the ventral precentral gyrus and two arrays in HCP-identified area 44 of the inferior frontal gyrus (considered part of Broca’s area). 1,155 days after the initial left hemisphere array implantation, four arrays were additionally implanted in the right hemisphere: two in HCP-identified area 6v of the right ventral precentral gyrus, and two in HCP-identified area 55b of the middle precentral gyrus. For left-hemisphere placement of arrays, see Willett et al., 2023 for details. Data are reported from post-implant days 30–149 and 1,185–1,304.

T15, a man with ALS, had four 64-channel microelectrode arrays placed in his left hemisphere: two in HCP-identified area 6v of ventral precentral gyrus, one in area 55b of middle precentral gyrus, and one in Brodmann area 4 (primary motor cortex); see ^4^ for details. T15 was 45–47 years old at the time of data collection, and the data reported are from post-implant days 25–636.

T16, a woman with pontine stroke, had four 64-channel microelectrode arrays placed in her left hemisphere: two in HCP-identified area 6d (hand knob), one in area 6v of ventral precentral gyrus, and one on the border of the premotor eye fields (PEF) and speech-related 55b; see ^13^ for details. T16 was 52–53 years old at the time of data collection, and the data reported are from post-implant days 99–596.

#### Functional MRI speech lateralization and array placement targeting

Participants T12, T15, and T16 underwent pre-operative anatomic and functional brain imaging for speech and language localization, surgical planning, and array placement targeting (see ^3,4,7,66^ for array location estimates and further details). See Extended Fig. 1 for the array location estimates of each participant.

#### Neural signal processing

Neural signals were recorded from the microelectrode arrays using the Neuroplex-E system (Blackrock Microsystems) and transmitted via a cable attached to a percutaneous connector. Signals were analog filtered (4th order Butterworth with cutoff frequencies 0.3–7.5 kHz), digitized at 30 kHz (250 nV resolution), and provided to a custom MATLAB Simulink or Python-based BRAND software^67^ for digital filtering and feature extraction. For T12 (up to 2024.07.09) and T16 (beginning from 2024.04.19), digital filtering began with a non-causal highpass filter (250 Hz cutoff) using a 4 millisecond delay. For T12 (beginning from 2024.04.19) and T15 (all session days), digital filtering instead began with a non-causal bandpass filter (250 Hz to 5 kHz cutoff) applied using a 1 millisecond delay. This non-causal zero-phase filtering was used to improve spike detection^68^. For all participants, linear regression referencing (LRR)^69^ was then applied to each array-group of 64 electrodes in order to reduce noise artifacts. Then, for all participants, two estimates of neural ensemble activity were computed for each electrode in 20 millisecond bins. Binned spike band power was computed by taking the sum of squared voltages at each time bin, and threshold crossing rate features were computed by counting the number of times that the filtered voltage time series crossed an amplitude threshold set at the standard deviation of the voltage signal multiplied by a constant factor (–3.5 for T12 and T16; –4.5 for T15). Threshold crossing rates and spike band power measure local spiking activity, and have been shown to yield similar decoding accuracies and neural population structure as sorted single neurons^70–73^.

#### Data collection rig

Digital signal processing and feature extraction were performed on a dedicated computer. For T12 sessions conducted prior to 2024.07.09, data was processed using Simulink Real-Time, and task software was implemented with the MATLAB Psychophysics Toolbox^74^. An additional Windows computer controlled the starting and stopping of tasks and interfaced with the Neuroplex-E system (Blackrock Microsystems, Salt Lake City, UT, USA). BRAND^67^, a framework for developing modular, Python-based neural data processing and task software, was used for all T12 sessions conducted from 2024.07.09 onwards, as well as for all T15 and T16.

#### Instructed delay paradigm for speech

All data collection sessions employed an instructed delay paradigm, with each trial consisting of a “delay” period followed by a “go” period. During the delay period, a sentence was displayed on the screen above a red square, allowing the participant time to read it and prepare to speak the sentence. After the delay period, the red square turned green with a chime to indicate the start of the go period. During this period, the sentence remained on the screen while the participant attempted to speak the sentence at their own pace. Once the participant had finished speaking the prompted sentence, progression to the next trial could be triggered by the participant pressing a button held in their lap (T12, T16), or selecting a button using an eye tracker mounted on the screen or using gesture decoding (T15). Each session was typically composed of multiple blocks of 40–50 trials. Between blocks, all participants were encouraged to rest as desired. All data collection sessions were performed at the participant’s place of residence.

During real-time evaluation, participant T12 attempted to produce vocalized speech to the best of her ability (although paralysis rendered the speech unintelligible). In sessions included in the offline dataset, participants either attempted vocalized speech or attempted mouthed speech (as in ^3^; refer to Table S1 of ^13^ for a detailed description of each speech mode), where the participant produced no audible sound but attempted to move the lips, tongue and jaw as though mouthing to someone across the room.

Each speech session began with a “diagnostic” block, where the participant was instructed to attempt to vocalize one of seven words or to “do nothing” for each trial (as in ^3^). The diagnostic block was used to calculate the spike threshold values and LRR coefficients that were applied to signal processing and feature extraction throughout the session for participants T12 and T16. For T15, the spike thresholds and LRR coefficients were re-calculated after each block.

For a summary of all data collection sessions, see Table 3.

#### Sentence selection

Sentences were sourced from different corpora covering a range of vocabularies and complexity. Sentences were primarily sourced from the Switchboard corpus^75^. The Switchboard corpus was also used for real-time evaluation. Some additional sentences were sourced from the OpenWebText2 corpus^76^ and the Harvard sentences^77^. A few sessions contained sentences consisting of a small 50-word vocabulary^1^, which was only used for training and not evaluation. In addition, for T15, a few sessions contained random orderings of words, or sentences constructed with words frequently used by T15 during his personal use of a speech BCI, both sourced from a 125,000-word vocabulary.

#### Decoder training data

In all analyses, we excluded features from arrays found to lack tuning to phonemes: arrays implanted in left hemisphere area 44 of participant T12^3^, and arrays implanted in areas 55b/PEF of participant T16^7^. Furthermore, all datasets were z-scored, and split into training (for training the base decoders), validation (for fine-tuning the LLM merger), and test sets (for evaluation). The validation and test sets were of roughly similar size.

For participant T12, the left hemisphere dataset was identical to that used in the Brain-to-Text ‘24 benchmark. To enable direct comparison of our results with the Brain-to-Text ‘24 benchmark, we used the same block-wise z-scoring procedure as in ^3^. For both the validation and test sets, a small set of blocks was randomly selected and held out from the dataset for evaluation.

For participant T15, the dataset was identical to that used in the Brain-to-Text ‘25 benchmark. As in ^4^, neural data for each trial was z-scored using a rolling window of 10 trials (mean and standard deviation were computed using the preceding 10 trials). A small set of blocks was randomly selected and held out from the dataset for evaluation, and then evenly split across the validation and test sets.

Participants T16 and T12 right hemisphere datasets were pre-processed similarly to the participant T15 dataset. Neural data for each trial was z-scored using a rolling window of 20 trials. For each session, the last 30% of trials in chronological order were held out for evaluation, and then evenly split across the validation and test sets. When less than 15 sentences were available for both the validation and test set, the entire session data was used as the training set.

### Online decoding with Participant T12

#### Recurrent neural network (RNN) architecture and training

The base decoder for online decoding was the same as that used in our prior work^3–5,7^. We used a 5-layer, stacked gated recurrent unit (GRU) RNN to map time series of neural features to sequences of phonemes.

To adapt to nonstationarities in neural features, a day-specific trainable input layer was added before the RNN. Connectionist temporal classification (CTC) loss^78^ was used with the Adam optimizer^79^ to train the RNN without requiring ground truth knowledge of when each phoneme was attempted.

Each RNN of the ensemble was initialized with random weights during training. To improve decoder stability, white noise and artificial constant offsets were applied to the input data during training, in addition to L2 regularization. Furthermore, to improve RNN performance, we included historical data from earlier sessions (post-implant days 30–149, when left hemisphere arrays yielded higher signal quality) in addition to newly collected data (from post-implant days 1185 up to the respective session day) during training. Since these earlier session data were collected before the right hemisphere arrays were implanted and thus had a smaller feature dimension, we zero-padded the missing right hemisphere features for these training examples, and applied a mask to prevent gradient flow to nonexistent features during training. A full list of RNN hyperparameters are available in Table 4.

#### RNN online recalibration

Each RNN was continuously recalibrated during evaluation to ensure decoding performance remained consistent throughout a session, as done in prior work^4,40^. At the beginning of a new session, each pre-trained RNN was loaded, and the most recent session’s day-specific input layer was duplicated. After each trial in the new session, the day-specific input layer and the weights of each base RNN were updated using the neural data and ground-truth sentences from each trial. To prevent overfitting to the current day’s data, each training batch contained 30% of historical data from previous sessions. A full list of RNN hyperparameters during online recalibration are available in Table 4.

#### n-gram model and language model rescoring

After each RNN emitted phoneme probabilities from neural activity, a 5-gram language model was used to predict a set of word-level sequences that best explain these probabilities while also being consistent with the statistics of the English language. The language model was implemented using a weighted finite state transducer (WFST) built using Kaldi^80^. The 5-gram language model uses a Viterbi (beam) search to combine the WFST graph’s transition probabilities with each RNN’s phoneme probability outputs, thereby inferring one or more likely word sequences and their associated likelihood scores. We used a pruned version of the original 125,000-word 5-gram model to accelerate real-time inference.

To provide feedback with partial predictions during real-time inference, the 5-gram model incrementally updated its word-level prediction during the trial as new phoneme probabilities were emitted from one arbitrarily chosen RNN. This partial prediction was updated on the screen in real-time.

After participant T12 indicated the end of her speech with a button press, the language models finalized their predictions. Given the full output probability sequence produced by each RNN, the 5-gram language model generated the 100 most likely word sequences and their corresponding scores. Subsequently, a decoder-only LLM (OPT-6.7B^81^) was used to rescore the candidate sequences. For each base decoder, candidate scores from the 5-gram language model were combined with those from the rescoring LLM using a weighted sum, and the candidate with the highest combined score was selected for each base decoder. The rescoring LLM weights were loaded using 4-bit quantization and computation was performed in 16 bits (bfloat16).

A full list of language model hyperparameters during training and inference are available in Table 4.

#### Merger LLM fine-tuning and inference

We used the GPT 4.1 mini model^51^ as the merger LLM, which was accessed through the OpenAI platform (OpenAI, San Francisco, CA, USA) for fine-tuning and inference. We used the prompt format introduced in ^28^, which includes both word and phoneme-level predictions from the base decoders. For both training and inference, following ^28^ we also appended the immediately preceding hypothesis set to the prompt as an in-context learning (ICL) example. During real-time operation, we evaluated three fine-tuning checkpoints and selected the output with the highest model-assigned probability. To minimize inference latency, each merger LLM checkpoint was called in parallel using asynchronous API calls, as supported by the OpenAI API.

#### Ensemble BCI system and network architecture

Our ensemble BCI system executed different processes across multiple computers (see Extended Fig. 2a for architecture diagram; Extended Figs. 3 and 4 for finite-state machine representations of processes). To enable inter-process communication (IPC) across these machines, we primarily used the BRAND^67^ software, which is designed to coordinate processes and exchange data between multiple machines via a Redis in-memory database.

##### Local decoder deployment and IPC

The local rig consisted of multiple computers, each supporting different system processes. These computers were connected via a local area network (LAN) for low-latency communication. One computer (running Ubuntu 22.04 LTS) hosted the Redis server, processed raw 30 kHz neural data, and extracted neural features. This machine was also capable of sending commands to other Redis client machines specifying processes and scripts to execute. Two additional computers (both running Ubuntu 22.04 LTS) were used to locally deploy one base decoder for low-latency feedback during closed-loop control. One computer was responsible for local RNN inference and recalibration, while the third computer performed decoding via the 5-gram language model and the rescoring LLM. A finite-state machine for the local decoder process is given in Extended Fig. 3.

##### Cloud architecture

With the exception of one locally deployed base decoder, the remaining base decoders of the ensemble BCI were distributed across 20 virtual machines on the Google Cloud platform (Google Cloud, Mountain View, CA, USA). Because each base decoder consisted of a cascade of subcomponents, including the RNN, 5-gram language model, and rescoring LLM, we deployed each of these subcomponents on separate cloud virtual machines. Therefore, nine virtual machines performed inference and recalibration of each RNN, and nine additional virtual machines executed LLM rescoring. The 5-gram language model was designed using Kaldi which natively supports multi-threading. A single shared instance, implemented on its own virtual machine, was used across the cloud-deployed base decoders.

##### Bidirectional rig-to-cloud communication

To enable bidirectional communication between the local rig and cloud infrastructure, we deployed an OpenVPN (OpenVPN Inc., Pleasanton, CA, USA) server on the cloud platform, and configured the local rig and cloud-based RNN virtual machines as its clients. This allowed us to stream neural data, task states, and commands from the local rig to the cloud virtual machines, and receive decoded outputs from the cloud in return via the common Redis database.

During real-time evaluation, a background thread was used to stream neural data and task states to the cloud virtual machines through Redis streams. After the end of a trial was indicated by the participant, each base decoder generated its predictions on cloud and streamed these predictions back to the local rig.

##### Intra-cloud communication

Routing messages through the local Redis server from the cloud virtual machine unavoidably introduces network delay. As a result, communication between the cloud virtual machines used WebSocket, instead of the locally hosted Redis server. The 5-gram language model and rescoring LLM virtual machines were each configured as a WebSocket server, and the corresponding RNN virtual machine was configured as their client. The RNN virtual machines were responsible for communicating with their downstream language model virtual machines via WebSocket, and with the local rig via Redis over the VPN gateway. Finite-state machines for the cloud-based processes are given in Extended Fig. 4.

#### Real-time evaluation session design

Each real-time evaluation session consisted of six to eight blocks of 40 trials. To reduce confounds due to early-session decoder miscalibration, the first one or two blocks were used as recalibration blocks. During these blocks, the decoded output was not displayed to the participant, but the decoder backend performed RNN recalibration in real-time. Closed-loop evaluation started after 900 to 1,800 gradient update steps for each base RNN, where training loss stabilized. This design allowed the RNNs to adapt to nonstationarities in neural data before closed-loop evaluation, and prevented poor early decoder performance influencing participant behavior during the subsequent closed-loop blocks. Closed-loop evaluation proceeded in alternating blocks of the single-decoder baseline and the ensemble.

#### Offline accuracy evaluation

To compare ensemble and baseline performance on identical sentences, while still using the decoder states generated during real-time operation (as displayed in Fig. 2c), we performed an offline simulation. During real-time evaluation, we configured single-decoder baseline blocks to still return final predictions from all base decoders in the background, although only the prediction from one base decoder was displayed to the participant. Offline, we retrospectively aggregated the saved base-decoder hypothesis set using the same merger LLM used during real-time evaluation. During ensemble blocks, the baseline decoder output was already available as one of the ten base decoders. Word error rates were subsequently computed for the offline single-decoder baseline and ensemble predictions, respectively.

#### Offline replay and optimization

To evaluate the viability of low-latency ensemble decoding, we separately optimized each component of the ensemble BCI. We then deployed nine base decoders on cloud and tested the viability of reducing cloud-invoked delay via an offline replay of the real-time evaluation sessions.

##### Relay network architecture

The original implementation independently streamed neural data from the local rig to each cloud RNN process over Redis, creating redundant local-to-cloud communication. We instead introduced a relay node colocated with the VPN server on the cloud platform. Neural data was streamed once from the local rig to this relay node and stored in a buffer, which was simultaneously streamed to each RNN virtual machine over WebSocket, using the high-throughput intra-cloud network.

##### RNN inference

To reduce RNN inference latency, the RNN inference function was pre-compiled as a dataflow graph using TensorFlow^82^. The computation graph was pre-computed at the start of the block, avoiding repeated graph retracing and compilation during inference. During offline replay, we used the same base RNN models as the real-time evaluation sessions.

##### RNN online recalibration

Since the RNN inference and online recalibration processes were co-located on the same machine, neural features, task state variables, and labels required for recalibration were maintained in a local shared memory buffer. The recalibration logic and hyperparameters were otherwise identical to that used during the real-time evaluation sessions.

##### Merger LLM inference

To accelerate merger LLM inference, we replaced remote API-based inference with local inference using efficient key-value cache management via vLLM^43^. We fine-tuned Qwen3-4B using 4-bit quantized low-rank adaptation^42^ with the Unsloth package^48^, using the same merger fine-tuning dataset as in the real-time evaluation. During inference, we evaluated the final three fine-tuning checkpoints and selected the output with the highest probability. The merger LLM was deployed locally.

To also reduce repeated computation across trials, we reused key-value tensors for the fixed instruction portion of the LLM prompt, so that only the trial-specific ensemble hypotheses needed to be newly processed on each inference call.

##### n-gram model and language model rescoring

To reduce n-gram decoding latency, we replaced the original Kaldi-based 5-gram model with an existing lightweight implementation using KenLM^64,83^. This software is available at: https://github.com/Neuroprosthetics-Lab/phoneme-to-words-lm. For language model rescoring, we used OPT-6.7B^81^ without 4-bit quantization, since quantization overhead can affect inference speed. Candidate sequences were then scored and selected using the same rescoring procedure as in the real-time system. Both 5-gram decoding and language model rescoring used the same parameters used during real-time evaluation sessions.

##### Simulation of real-time evaluation sessions

To demonstrate the robustness of our method, we performed offline simulation under slightly more suboptimal conditions than those during real-time evaluation. We measured the participant’s home network connection to the VPN server using iPerf3, in 1 second reporting intervals over 300 seconds. We then throttled the VPN link to 20 Mbps, which was the lowest throughput observed over the 300 second measuring period. We also ensured that the average round trip time (measured over 100 pings) during offline simulation (67 milliseconds) was at least equal to or worse than the participant’s home network (52 milliseconds).

Next, to replay the real-time evaluation session data, neural data and end-of-speech events were emitted from the local rig according to the timestamps logged during the original real-time evaluation. Using the same hardware and decoders used during real-time evaluation, we measured the duration of time until the final prediction was produced after the end of speech, and completion times for each decoding step.

### Decoder performance metrics

We evaluated the decoder using four metrics: word error rate, words per minute, latency, and hardware utilization.

#### Word error rate

Word error rate was defined as the edit distance between the decoded sequence of words and the cue sentence. To compute “aggregate” word error rates, which are reported in this study, we divided the total number of errors by the total number of words across all sentences.

#### Words per minute

Words per minute was defined as the number of words spoken divided by the total duration of speaking time. Speaking time for each trial was measured as the time from the go cue until the participant pushed the button to signal the end of speech. The aggregate words per minute was also calculated by dividing the total number of words by the total speaking time across all sentences.

Confidence intervals were computed via bootstrap resampling 10,000 times over individual trials. Statistical significance tests comparing the baseline and the ensemble were performed using a paired permutation test over matched utterances (10,000 permutations). The predicted labels by the baseline and the ensemble were randomly swapped within each utterance pair, and the total word error rate difference was recomputed for each permutation. Two-sided *p*-values were calculated as the fraction of permutations whose absolute difference was at least as large as the observed absolute difference.

#### Inference latency

Third, latency to final prediction shown in Fig. 3a was measured as the duration of time for producing a final prediction after the participant pushed the button to signal the end of speech. Latency to final prediction in Fig. 3c was measured in the same way, except during an offline replay that replicated in-session conditions such as network quality, neural data, and button press timings.

Cumulative latency shown in Fig. 3b was measured as the duration of time until each decoding step was fully completed. For cumulative completion times of the cloud-based decoding steps, we used the completion time of the slowest base decoder during each trial. To illustrate step-wise completion times for a typical trial as tail events can make interpretation of the results difficult, only latencies up to the 90th percentile of trials for each decoding step were visualized. The full distribution of step-wise completion times were visualized in Fig. 3d.

Step-wise latency shown in Fig. 3b and d were measured as the time required for each decoding stage (neural data readout, RNN phoneme decoding, language model decoding, and hypothesis aggregation) to complete. For decoding stages performed by the base decoders, step-wise latency statistics for the slowest base decoder were plotted.

For Fig. 3 subpanels, two-sided *p*-values were calculated as the fraction of permutations whose absolute difference was at least as large as the observed absolute difference. Furthermore, for visualization purposes, the violin plots in Fig. 3 were smoothed using a Gaussian kernel density estimate with a standard deviation approximately equal to 25% of the data’s standard deviation.

#### Hardware utilization

Finally, we evaluated hardware utilization during offline replay of the optimized ensemble BCI, as summarized in Table 2, by monitoring the equivalent vCPU cores, peak system memory usage, and peak GPU memory usage during each trial’s decoding window. Equivalent vCPU cores were defined as the process CPU time accumulated during the decoding window, summed across all process threads, divided by the corresponding wall-clock elapsed time. Thus, a value of 1.0 indicates use of one full vCPU core, whereas values greater than 1.0 indicate parallel CPU use across multiple cores. System memory usage and GPU memory usage were defined as the memory used on the corresponding hardware, sampled every 10 ms during each trial’s decoding window. For each hardware metric, we computed the peak value observed within each trial’s decoding window and reported the highest observed value across all trials in Table 2.

The decoding window for each trial was defined as the interval within a trial during which a given machine was actively performing the trial-specific decoding computation. This was to ensure that utilization estimates accurately reflected active computational load of the decoding task, rather than being diluted by idle time, network waiting, or computation occurring on other machines. For each RNN decoder, the decoding window extended from when the last neural bin arrived to when phoneme probabilities were computed; for the 5-gram language model, from when decoding began on the first available RNN phoneme probability sequence to when the final 5-gram hypothesis set was produced; for each rescoring LLM, from when the 5-gram hypothesis set was received to when the rescored prediction was produced; and for the merger LLM, from when all candidate predictions were received to when the merger prediction was produced.

### Offline decoding and analysis

#### Base decoder

The offline base decoder architecture was similar to the online decoder architecture, with a few modifications to improve offline decoding accuracy. Firstly, to enable direct comparison with results of the Brain-to-Text benchmarks, we used bidirectional RNNs, and optionally appended an expanded classification head (two blocks each consisting of layer normalization, dropout, a fully connected layer, and a nonlinear activation). This minimally modified architecture was demonstrated in one of the entries for the Brain-to-Text ‘24 benchmark to improve accuracy^26^. RNNs were trained using either the Adam^79^ or AdamW^84^ optimizer under a CTC loss objective^78^. Each RNN’s weights were randomly initialized during training.

For RNNs evaluated on T12 left + right hemisphere data, initial training included earlier sessions with only left hemisphere features by using feature zero-padding and gradient masking, similar to the online system. To improve accuracy, the RNNs were then fine-tuned only on sessions in which right hemisphere features were available.

The 5-gram language model decoding and LLM rescoring were also performed using the same method as the online system. For offline decoding, the unpruned full 125,000-word 5-gram language model and full precision (32-bits) OPT-6.7B model were used. Language model hyperparameters (see Table 5) were based on the values reported in prior studies for each respective participant^3,4,7^.

#### Merger LLM fine-tuning and inference

After RNN training, we generated hypothesis sets from the validation set and used this data to fine-tune the merger LLM (Qwen3-32B^44^). Fine-tuning and inference were performed using 4-bit quantized low-rank adaptation^42^ via the Unsloth package^48^. Prompt construction was performed in the same way as the online system. In cases of text degeneration (detected as abnormal length of five standard deviations above the mean, repetition of the prompt, or production of non-valid characters), we replaced the output with a base decoder prediction. Text degeneration was very rare (0.004% across all merger LLM inference calls performed in this paper), with the highest per-condition observed rate (under the 50-sentence fine-tuning condition) still being 0.07%.

All merger LLM hyperparameters can be found in Table 6. To choose the hyperparameters, we re-partitioned the original training set of an arbitrarily selected dataset (T12 left hemisphere) into a training set (used for RNN training) and a validation set (used for LLM fine-tuning). The original validation set was used to evaluate the merger LLM’s word error rate, and the original test set (used to report the ensemble results in this paper) was excluded in hyperparameter selection.

As it can be computationally intensive to fine-tune LLMs across a grid of hyperparameters, we performed 70 fine-tuning runs with Bayesian optimization to iteratively determine which hyperparameter configurations were most likely to minimize word error rate (based on previous hyperparameters and their performances). The choice of hyperparameters that yielded the lowest word error rate was selected. To prevent over-optimization to a specific dataset, and since LLM hyperparameter sweeps can be computationally costly in realistic deployment settings, the same merger LLM hyperparameters were used across all participants.

#### Pseudoensembling

Formally, pseudoensembling was performed by applying a noise process to a trial neural activity, represented as a discrete-time matrix 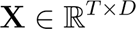 where *T* is the number of time samples and *D* is the number of features. The noise injection process generated a total of *N* - 1perturbed versions 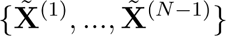, each defined by

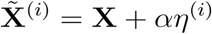

where each noise template time-series 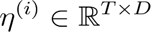 was sampled from an discrete-time zero-mean multivariate Gaussian distribution 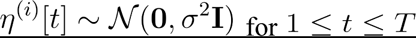. Each noise template was scaled by a positive scalar 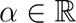. The original and perturbed neural activities were then each passed through an identical decoding pipeline (consisting of a single RNN, 5-gram model, and re-scoring LLM), represented as 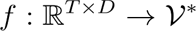 where *V\**represents all finite-length sequences that can be formed with the vocabulary *V*.

A hypothesis set of size *N* was defined as

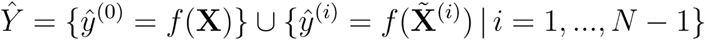

#### A fine-tuned merger LLM was used in the same way as full ensembling to aggregate the hypothesis set

The noise perturbation strength α was tuned for each session day to best reproduce the disagreement level between hypotheses used in the full ensemble. To measure the ensemble disagreement level, we first independently trained two base decoders, and measured the word error rate between the two hypotheses (measured as word error rate, treating one prediction as reference) for each session day. Then, we swept across 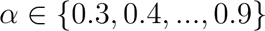 and generated two independently perturbed versions of neural data. We passed both versions through a single base decoder, and computed the resulting word-level disagreement between the two predictions for each session day. Finally, we selected the session day-specific perturbation strength as the α whose pseudoensemble disagreement was closest to the full ensemble disagreement in absolute difference.

#### Evaluation under various design considerations

We evaluated the full ensemble under three design considerations: hypothesis aggregation method, fine-tuning dataset size, and ensemble size.

##### Hypothesis aggregation method

Different hypothesis aggregation methods were compared: ROVER, GPT-family models (GPT 4.1, 4.1 mini, and 4.1 nano), Llama 3.1 8B, and Qwen3-32B (Fig. 5).

Our implementation of ROVER followed ^33^. First, dynamic programming was used to compute the minimum-edit alignment between two hypotheses. This alignment was used to form an initial slot-based confusion network where each aligned position stored alternative word hypotheses and their cumulative weights, with epsilon representing deletions. Additional hypotheses were then incorporated incrementally by aligning each new word sequence to the confusion network. After constructing the confusion network, the final consensus transcript was generated by taking a majority vote across each aligned word position.

LLM-based hypothesis aggregation methods were performed using separate procedures depending on the model family of the LLM. Fine-tuning and inference for the GPT-family was performed via the OpenAI platform (OpenAI, San Francisco, CA, USA). For both Llama 3.1 8B and Qwen3-32B, fine-tuning and inference were performed via 4-bit quantized low-rank adaptation^42^ using the Unsloth package^48^. Default hyperparameters were used for Llama 3.1 8B (Table 6).

Each aggregation method was evaluated across 3 independent runs by re-training the base decoders and aggregation method (if necessary) from distinct randomly initialized weights. The mean word error rate for each aggregation method is reported along with 95% confidence intervals in Fig. 5.

##### Dataset size

To evaluate the effect of merger LLM fine-tuning set size on accuracy, we separately fine-tuned the merger LLM on randomly subsampled fine-tuning sets of approximately 20, 50, 100, 200, and 400 samples. For participants with sufficient data, we also evaluated larger training set sizes. Each dataset size was evaluated across 10 independent random subsampling runs. The mean word error rate for each dataset size was reported along with 95% confidence intervals in Fig. 6a–d.

##### Ensemble size

Ensemble size was defined as the number of base decoders that were used to produce the hypothesis set. To evaluate the effect of ensemble size on accuracy, we separately trained the RNNs and fine-tuned merger LLM on varying ensemble sizes of 2, 5, 8, 10, and 20. Each ensemble size was evaluated across 5 independent runs, using distinct, non-overlapping sets of random weight initialization for each run during training. The mean word error rate for each ensemble size was reported along with 95% confidence intervals in Fig. 6e–h.

#### Literature survey

For Fig. 1c and d, we surveyed published and publicly available results in the Brain-to-Text ‘24 benchmark. For Fig. 1d, we compared single-decoder and ensemble word error rates evaluated on the benchmark. Only ensembles using ten base decoders are shown to assess whether ensemble gains can be observed across different baseline word error rate regimes, while avoiding differences in ensemble size as a confounding factor. When published or benchmark-reported values were available^28–31^, we plotted the best-performing word error rates reported by the original authors. For “LightBeam^30^,” because individual baseline word error rates were not available, we used the reported average baseline word error rate. For single-decoder methods with publicly released code but without reported ensemble performance (“Baseline GRU^3^” and “Linderman GRU^26^”), we generated ensemble results using one run of ten independently initialized base decoders and a fine-tuned merger LLM (GPT 4o-mini^85^).

## Extended figures

**Extended Fig. 1.**
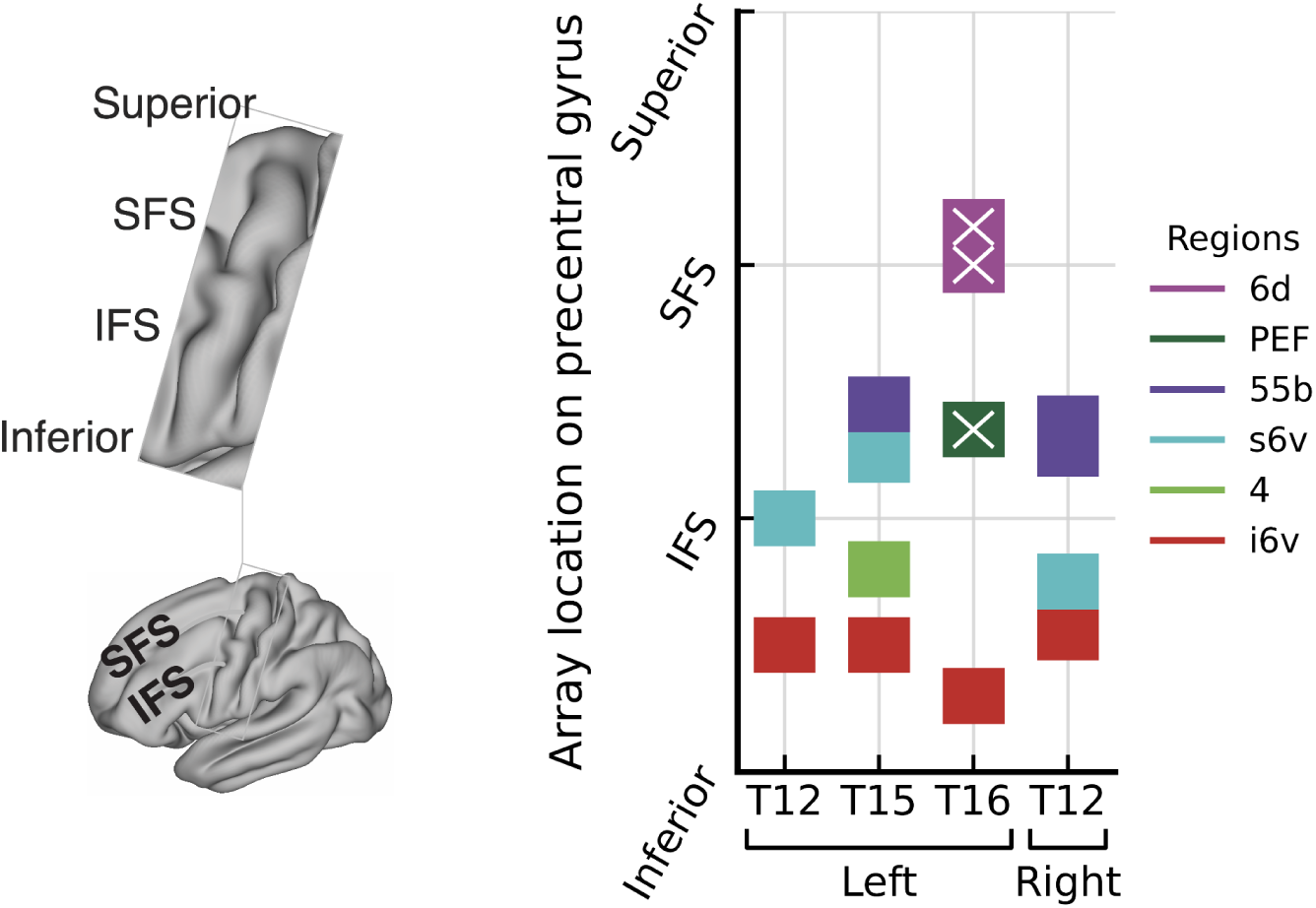
Approximate array locations of participants on precentral gyrus. Zoomed-in view of the precentral gyrus of an averaged brain (left) and color-coded approximate array locations (right). On the y-axis, IFS and SFS represent the inferior and superior frontal sulcus, respectively. T16 arrays with little tuning to speech motor movements (and therefore were not used for analysis) are crossed out.

**Extended Fig. 2.**
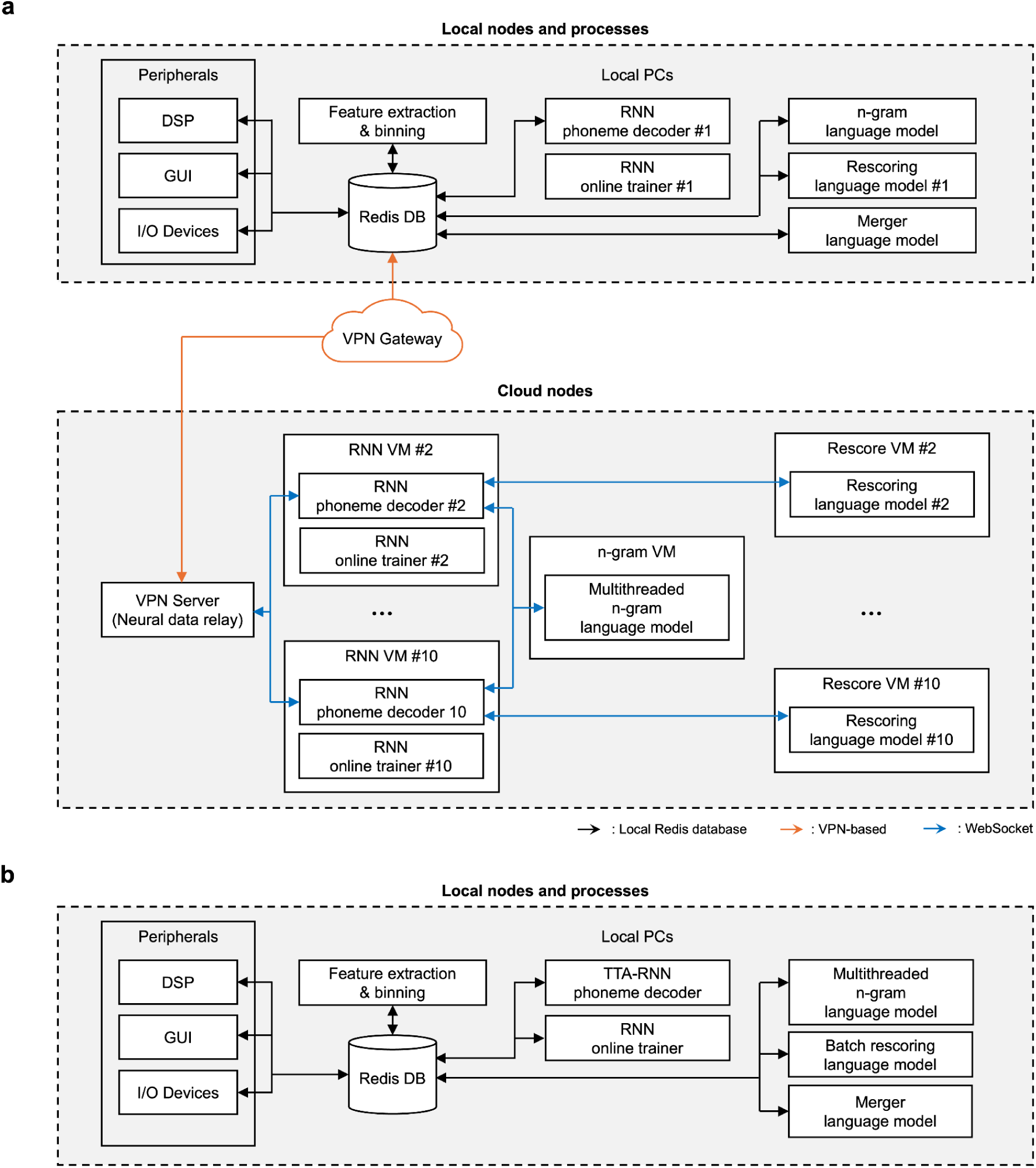
System and networking diagrams for ensemble speech BCIs. **a,** System and networking diagram for the ensemble speech BCI. **b,** Prospective system and networking diagram for a pseudoensembling speech BCI.

**Extended Fig. 3.**
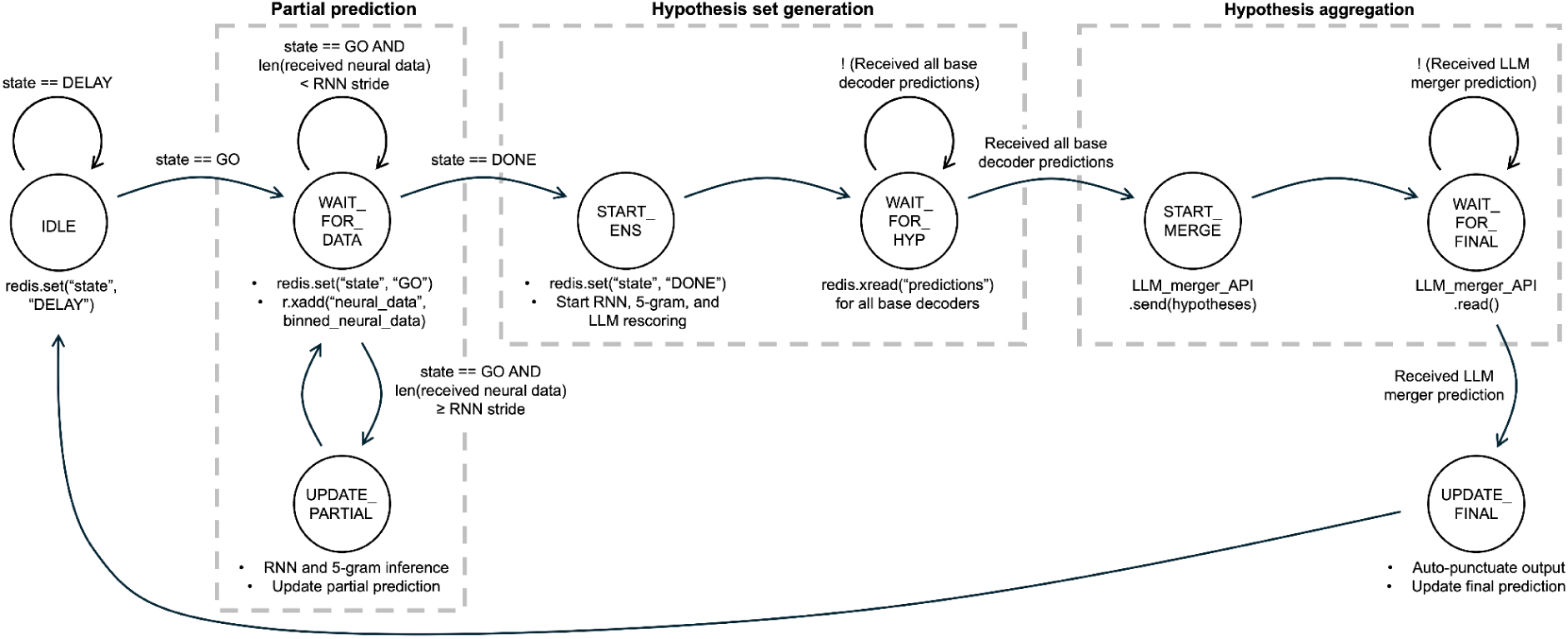
Operational finite-state machine (FSM) diagram for the locally deployed decoder. While the participant is speaking, the locally deployed decoder iteratively generates a partial prediction. After the participant is done speaking, the decoder initiates ensemble decoding on the cloud infrastructure, and updates the final prediction. This modular design allows decoder subcomponents to be easily added, substituted, or removed with minimal changes depending on the BCI architecture.

**Extended Fig. 4.**
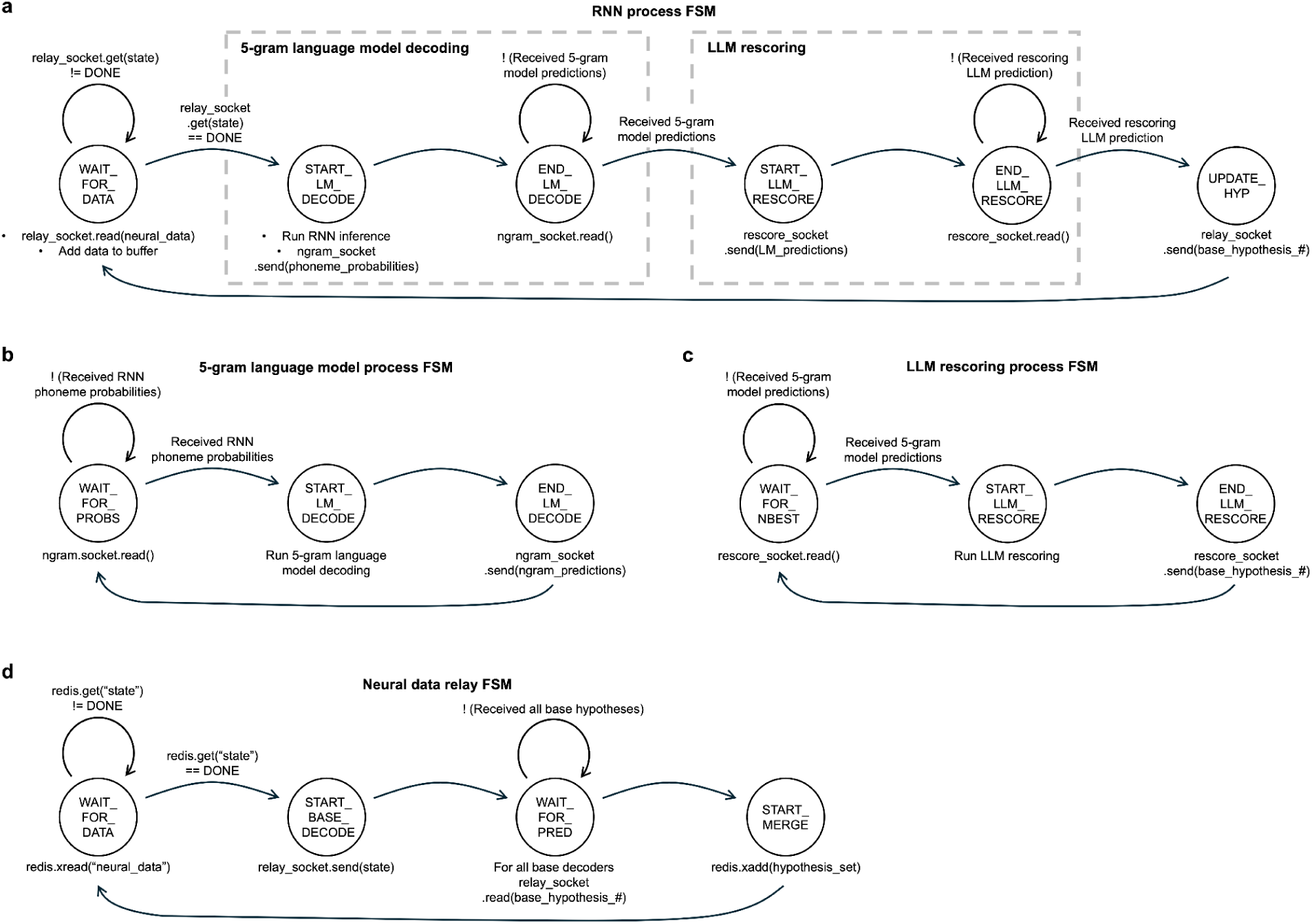
Operational finite-state machine (FSM) diagrams for the cloud-deployed decoder components. **a,** FSM for the cloud-deployed RNN process. **b,** Same as in **a,** but for the 5-gram language model process. The single-thread version is shown. **c,** Same as in **a,** but for the LLM rescoring process. **d,** same as in **a,** but for the neural data relay.

